# Cytosolic Ribosomal Protein Haploinsufficiency affects Mitochondrial Morphology and Respiration

**DOI:** 10.1101/2024.04.16.589775

**Authors:** Agustian Surya, Blythe Marie Bolton, Reed Rothe, Raquel Mejia-Trujillo, Qiuxia Zhao, Amanda Leonita, Yue Liu, Rekha Rangan, Yasash Gorusu, Pamela Nguyen, Can Cenik, Elif Sarinay Cenik

## Abstract

The interplay between ribosomal protein composition and mitochondrial function is essential for sustaining energy homeostasis. Precise stoichiometric production of ribosomal proteins is crucial to maximize protein synthesis efficiency while reducing the energy costs to the cell. However, the impact of this balance on mitochondrial ATP generation, morphology and function remains unclear. Particularly, the loss of a single copy ribosomal protein gene is observed in Mendelian disorders like Diamond Blackfan Anemia and is common in somatic tumors, yet the implications of this imbalance on mitochondrial function and energy dynamics are still unclear. In this study, we investigated the impact of haploinsufficiency for four ribosomal protein genes implicated in ribosomopathy disorders (*rps-10, rpl-5, rpl-33, rps-23*) in *Caenorhabditis elegans* and corresponding reductions in human lymphoblast cells. Our findings uncover significant, albeit variably penetrant, mitochondrial morphological differences across these mutants, alongside an upregulation of glutathione transferases, and SKN-1 dependent increase in oxidative stress resistance, indicative of increased ROS production. Specifically, loss of a single copy of *rps-10* in *C. elegans* led to decreased mitochondrial activity, characterized by lower energy levels and reduced oxygen consumption. A similar reduction in mitochondrial activity and energy levels was observed in human leukemia cells with a 50% reduction in *RPS10* transcript levels. Importantly, we also observed alterations in the translation efficiency of nuclear and mitochondrial electron transport chain components in response to reductions in ribosomal protein genes’ expression in both *C. elegans* and human cells. This suggests a conserved mechanism whereby the synthesis of components vital for mitochondrial function are adjusted in the face of compromised ribosomal machinery. Finally, mitochondrial membrane and cytosolic ribosomal components exhibited significant covariation at the RNA and translation efficiency level in lymphoblastoid cells across a diverse group of individuals, emphasizing the interplay between the protein synthesis machinery and mitochondrial energy production. By uncovering the impact of ribosomal protein haploinsufficiency on the translation efficiency of electron transport chain components, mitochondrial physiology, and the adaptive stress responses, we provide evidence for an evolutionarily conserved strategy to safeguard cellular functionality under genetic stress.

## INTRODUCTION

The coordinated biogenesis of ∼79 ribosomal proteins (RPs) is crucial for cellular health and development. In humans, the haploinsufficiency or point mutations of ribosomal protein genes leads to a range of ribosomopathies^1^, including Diamond-Blackfan Anemia (DBA)^2^, and has been linked to an increased susceptibility to certain cancers, such as Myelodysplastic Syndromes and Acute Myeloid Leukemia^3,4^. With its hallmark features of hematological dysfunction and an increased risk of malignancies, DBA exemplifies the systemic consequences of ribosomal protein deficits^5,6^.

DBA and other ribosomopathies are rare genetic disorders. However, hemizygous losses of ribosomal protein genes are frequently observed (∼40%) in tumors^7,8^ and impact cellular proliferation and oncogenesis^9–11^. Specifically, ribosomal protein mutations are associated with higher mutational load in T-cell acute lymphoblastic leukemia(T-ALL) patients^12,13^. Hemizygous deletion of *RPL5* occurs in 11-34% of multiple tumor types, and reduced expression of this gene is correlated with poor survival in glioblastoma and breast cancer^14^. Conversely, overexpression of RPL15 and RPL28 lead to increased metastatic growth^15,16^. The phenotypes associated with these genetic disruptions allude to roles that these proteins play beyond protein synthesis.

Considering the significant energy demands of ribosome biogenesis^17^, mitochondrial function and ribosome production are interconnected to ensure optimal cellular energy equilibrium. A reciprocal connection between ribosomal and mitochondrial DNA copy number is observed across individuals^18^. Moreover, mitochondrial dysfunction leads to retrograde signaling that alters accumulation of extra chromosomal ribosomal DNA circles^19^. One potential mechanistic link between these processes is RNAse MRP, which is involved both in the processing of ribosomal RNA in the nucleolus, and in priming DNA replication in mitochondria^20–24^.

Other observations supporting the connection between ribosome biogenesis and mitochondrial function include: (i) Translation of mitochondrial transcripts are reduced and mitochondrial structure and oxygen consumption are altered in response to the deletion of ribosome biogenesis factor, Bud23 in mouse cardiomyocytes^25^. (ii) Yeast *Asc1* (*RACK1* ortholog) mutants, exhibit reduced translation of cytosolic and mitochondrial ribosome transcripts and lower fitness in a non-fermentable carbon source suggesting decreased mitochondrial activity^26^. (iii) The inhibition of ribosomal RNA synthesis through the depletion of the RNA polymerase I component, RPOA-2, in *C. elegans* results in a significant decrease in mitochondrial ribosomal proteins without affecting their transcript levels^27^. These observations suggest that altering ribosome biogenesis could alter mitochondrial components or function across different species.

Reciprocal to the evidence provided, biogenesis of cytosolic ribosomes also requires functional mitochondria. For instance, Rli1p, a protein carrying Fe/S clusters and thus requiring mitochondrial protein machinery, is associated with ribosomes and Hcr1p, which is involved in 20S pre-rRNA processing and translation initiation^28^.

Interestingly, a notable parallel has been observed between DBA and Pearson syndrome, which results from mitochondrial DNA losses^29^. Their symptoms are strikingly similar; in one instance, ∼5% of patients initially diagnosed with DBA were found to have significant mitochondrial DNA loss, leading to a reclassification as Pearson syndrome^30^. Similarly, expression analysis within a large family carrying a single-copy SNP variant in RPL11 suggested altered mitochondrial expression, indicating that coordination between mitochondria and ribosomes may be disrupted upon single-copy loss of ribosomal protein genes^31^.

Despite the known links between ribosome biogenesis and mitochondrial morphology, the ways in which mitochondrial function and oxidative stress relate to ribosomal protein (RP) haploinsufficiency have yet to be explored. Here we examine the effects of single copy loss for four ribosomal protein genes (*rps-10, rpl-5, rpl-33, rps-23*) in *Caenorhabditis elegans*, along with corresponding reductions in human lymphoblast cells. Our investigations revealed significant mitochondrial morphological alterations across these mutants and a conserved mechanism that coordinates the synthesis of mitochondrial components in response to compromised ribosomal machinery. Notably, a reduction in the cytoplasmically assembled RPS-10 in *C.elegans (rps-10(0)/*+ mutant), exhibited altered mitochondrial function and reduced cellular energy—a phenomenon mirrored by a 50% reduction in RPS10 abundance in human cells. These observations are further supported by significant expression covariation between mitochondrial membrane components and ribosomal proteins across lymphoblastoid cells derived from a diverse group of individuals, suggesting an adaptive conserved mechanism of mitochondrial function in response to ribosomal expression alterations.

## RESULTS

### Developmental and Physiological Consequences of Ribosomal Protein Gene Haploinsufficiency in *Caenorhabditis elegans*

We sought to determine the impact of haploinsufficiency of RP genes in *C. elegans*, and focused on the single-copy losses of two large subunit ribosomal proteins, *rpl-5 and rpl-33,* along with two small subunit RPs, *rps-10* and *rps-23* ^32^. We prioritized these four RP genes due to their involvement in human ribosomopathies, with *rpl-5*, *rpl-33*, and *rps-10* relating to DBA^33^, and *rps-23* relating to microcephaly and intellectual disability without the blood phenotypes^1^. Moreover, the protein products of these RP genes are incorporated into nascent ribosomes at different stages (nucleolar, nuclear and cytoplasmic)^34^. We observed developmental delays across these RP haploinsufficient mutants compared to wild-type counterparts (**Figures 1A,1B**, and **S1A**). Protein levels were evaluated using semi-quantitative proteomics against stage-matched controls, revealing reductions of approximately 50% for RPL-33 and RPS-23, 25% for RPL-5, and 10% for RPS-10 (**Figure 1C**, **Data S1**). The developmental delays observed in haploinsufficient strains, as compared to their time-matched controls, were found to generally correlate with the degree of protein reduction resulting from the loss of a single copy (**Figure 1C**). These findings suggest a correlation between the extent of protein level reduction and the timing of developmental processes.

**Figure 1:**
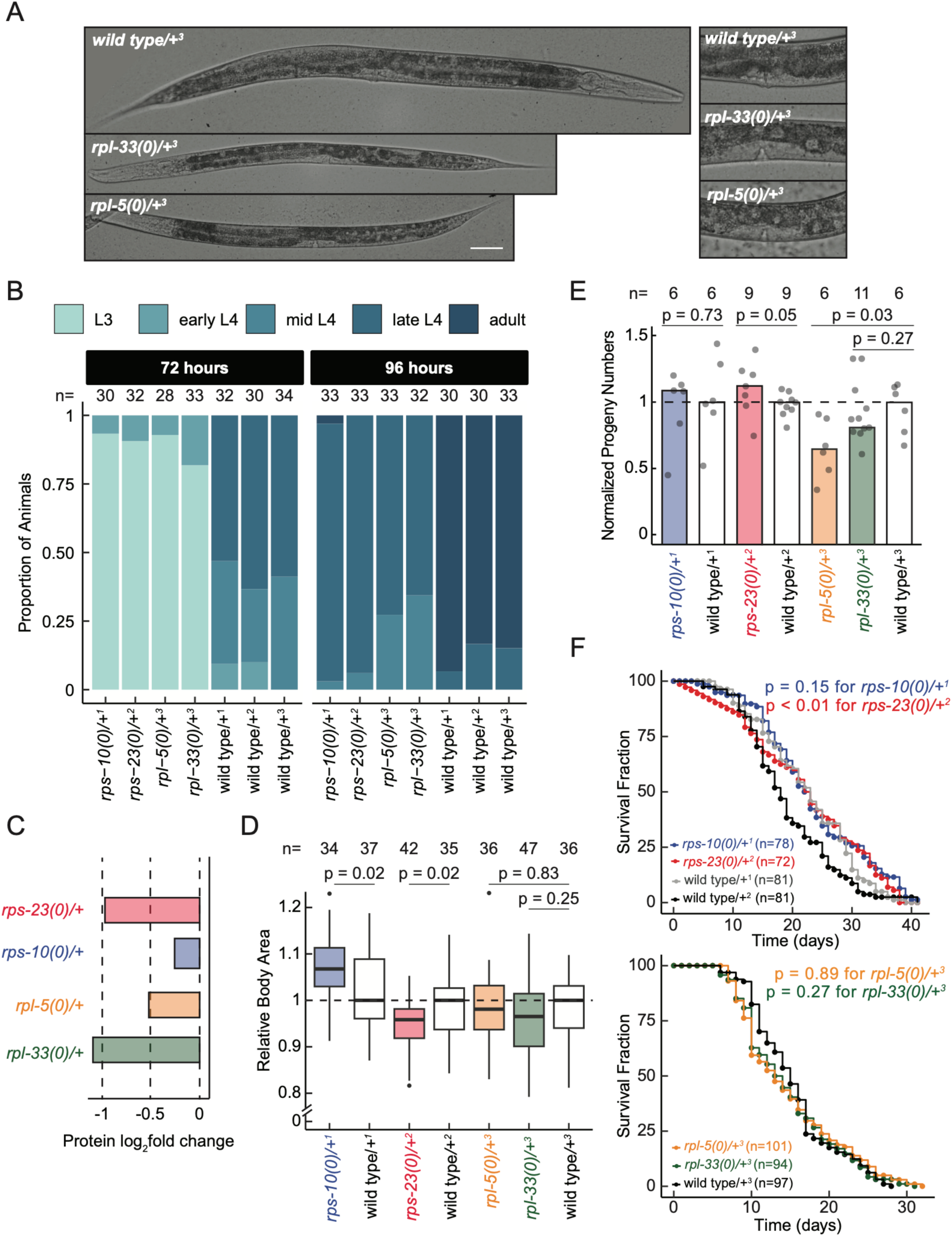
Ribosomal Protein Haploinsufficiency Results in Developmental Delays and Variable Body Size Without Affecting Lifespan in *C. elegans*. (**A**) The development of single copy large subunit ribosomal protein (RP) mutants alongside their wild-type counterparts after 96 hours of incubation from embryo at 16℃. Images show approximately one day of growth delay (left) and differences in vulval development (right), with a scale bar of 50 µm. Differential interference contrast images were taken with a 20X objective. (**B**) The developmental stage of animals after 72 and 96 hours of incubation from embryo at 16℃ were determined and plotted, with a stacked bar chart indicating the relative percentage of larvae at each stage. (**C**) Relative ribosomal protein levels quantified in haploinsufficient RP mutants using semi-quantitative proteomics of stage matched animals, with relative protein expression predicted by Differential Expression Proteomics (DEP package, R). (**D**) Body area of stage-matched animals (at late L4 stage), relative to the median body area of their respective controls were plotted. Differences in the mean relative body area of each mutant, compared to its respective control, were quantified using ImageJ and statistical significance was determined using an independent Student’s *t*-test. (**E**) Brood size of each haploinsufficient RP mutant, normalized to their respective wild-type controls, were plotted. Each dot represents an individual animal. Differences in the mean normalized brood size of each mutant, compared to its respective control, were analyzed using an independent Student’s *t*-test. (**F**) Lifespan of small subunit RP mutants (top) and large subunit RP mutants (bottom) alongside their respective wild-type controls are shown. Lifespan data are presented in a survival plot, with statistical analysis conducted using the Kaplan-Meier test and Bonferroni correction for multiple comparisons. Superscript numbers denote the specific wild type balancer chromosomes, and are used to compare between an RP mutant and the wild-type counterpart. Balancer chromosomes are denoted as follows: +^1^ = *tmC20*, +^2^ = *tmC5*, +3 = *mIn1*, All experiments were performed in at least three biological replicates, and the animals were grown at 16°C. In panels (C, D, E and F), RP mutants are color-coded for clarity: *rpl-33(0)/*+^3^ in green, *rpl-5(0)/*+^3^ in orange, *rps-10(0)*/+^1^ in blue, and *rps-23(0)/*+^2^ in red.

Similar to previous reports of reduced body size in RP mutants in other species^35,36^, our examination revealed that, when given sufficient time to reach the same developmental stage, *C. elegans* RP haploinsufficient mutants were slightly smaller in body size than wild-type controls with one exception (**Figure 1D**). Specifically, we observed that *rps-10(0)/*+ animals were slightly larger in body size compared to their stage-matched controls (**Figure 1D**, p=0.02, independent Student’s *t*-test). The increased body size could be associated with increased cell volume, altered cytoskeletal dynamics or metabolism due to changes in signaling pathways such as TGF-β, MAPK or cGMP^37–39^. These results are also reminiscent of the larger wing sizes and wing discs observed in *Drosophila* following the single copy loss of *RpL38* and *RpL5*^40^. However, why single copy loss of certain RP genes leads to increased organ or body growth remains unclear.

We further conducted fecundity assays to assess impacts on reproduction. With the exception of *rpl-5(0)/*+ animals, which exhibited a significant reduction in progeny size (∼25% reduction; p=0.03, independent Student’s *t*-test), the progeny sizes of all other RP haploinsufficient mutants were similar to those of the controls (**Figure 1E**). The onset of peak fertility was delayed in all mutants except for *rps-10(0)+/* animals (**Figure S1B**). Additionally, RP mutants remained fertile for extended periods, thereby compensating for the overall progeny size, with the exception of *rpl-5(0)/*+ mutants. Reduced fertility was observed in *Drosophila minute* mutants, characterized by the lack of a single copy of an RP gene^35^. The observation of similar brood sizes in the majority of RP strains in *C. elegans* suggests involvement of compensatory mechanisms within the germline, including a pattern of delayed but prolonged fertility.

Finally, we investigated lifespan in *C. elegans* RP mutants. Our lifespan analysis did not reveal significant differences for the majority of RP haploinsufficient mutants compared to wild-type controls, regardless of treatment with the egg-laying inhibitor Fluorodeoxyuridine (FuDR). The only exception was *rps-23(0)/*+ mutants that displayed a modest but significant extension of lifespan only in the absence of FuDR ( **Figure 1F**, **S1C**, p =0.007 for *rps-23(0)/*+ mutants, Kaplan-Meier test). Reduced protein translation or the knockdown of RP genes are typically linked to increased lifespan^41–46^. However, our results suggest that haploinsufficiency for single RP genes negate the typical lifespan extension benefits associated with decreased protein synthesis due to imbalances in RP expression and the associated stress. Taken together, our phenotypic characterization of the haploinsufficiency of RP genes in *C. elegans* reveals a range of developmental and physiological consequences that broadly mimic those observed in other model organisms such as *Drosophila* and mice^35,36^.

### Adaptive Cellular Responses to Ribosomal Protein Loss Highlights SKN-1 Dependent Enhanced Oxidative Stress Resistance

To understand the cellular mechanisms triggered by single copy losses of ribosomal protein genes in *C. elegans,* we performed RNA sequencing (RNA-seq) on stage-matched mutant and control animals at the L4 stage (**Data S2**). This analysis, which included stage-matched controls to mitigate any developmental delay effects, revealed a uniform gene expression response across all mutants. This response was characterized by overexpression of ribosomal machinery, glutathione transferase activity, and genes involved in innate immunity and stress responses, indicating a systemic adaptation to ribosomal protein loss (**Figure 2A**, top, significantly enriched gene annotation [GO] categories provided in **Data S3**). Conversely, genes related to mitochondrial activity, fatty acid biosynthesis, cell polarity, and amino acid metabolism were significantly underexpressed (**Figure 2A**, bottom, **Data S3**), suggesting a reprogramming of cellular metabolism in response to ribosomal protein haploinsufficiency.

Given the pronounced overexpression of glutathione transferase (*gst*) genes (∼2.8 fold enrichment, p<0.001, gene ontology (GO) enrichment), we assessed the expression patterns of *gst* genes across all RP mutants (**Figure 2B**). The general overexpression signature aligns with previously established links between glutathione transferase activity and oxidative stress resistance^47,48^, prompting us to assess the mutants’ resilience to oxidative stress.

**Figure 2:**
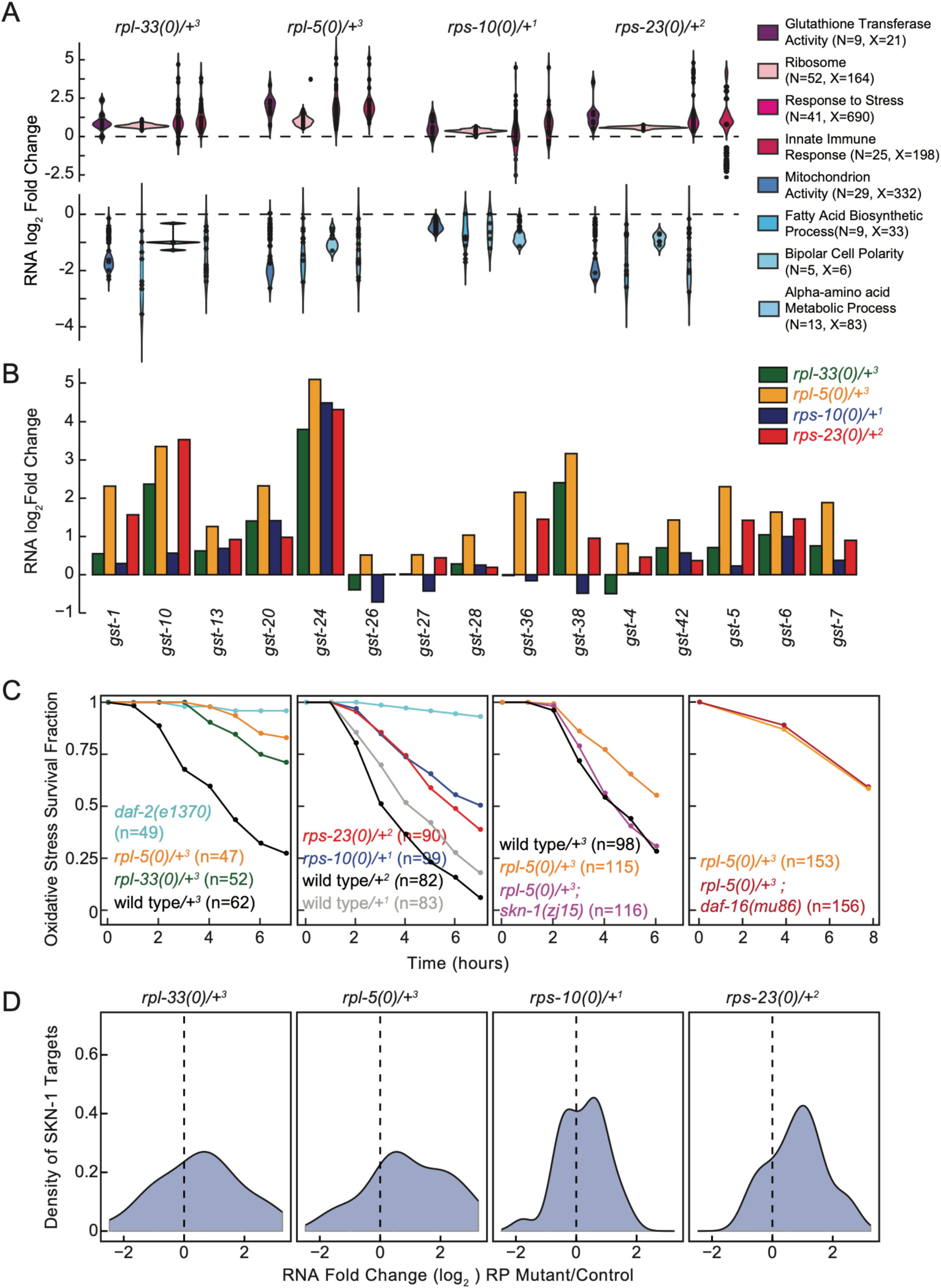
Adaptive Cellular Responses to Ribosomal Protein Loss Highlights SKN-1 Dependent Enhanced Oxidative Stress Resistance. **(A)** Investigation of significantly differentially expressed genes at the RNA level across all haploinsufficient RP mutants, with a focus on significant gene annotation–GO– enrichments are shown. Representative GO categories for overexpression (top) and underexpression (bottom) were plotted. The Y-axis displays predicted RNA-seq log_2_ fold changes. Gene Ontology (GO) categories are annotated on the side, where ‘N’ denotes the number of differentially expressed genes within a category, and ‘X’ represents the total number of genes in that category. Each dot signifies an individual gene identified as underexpressed or overexpressed, with their distribution across the categories visualized through violin plots. **(B)** RNA expression predictions of glutathione transferase genes annotated in the *C. elegans* genome were plotted across all mutants compared to their controls. **(C)** An acute time course of oxidative stress survival was plotted for large subunit RP mutants (first plot, *rpl-5(0)/*+ and *rpl-33(0)/*+) and small subunit RP mutants (second plot, *rps-23(0)/*+ and *rps-10(0)/*+), alongside wildtype control strains and *daf-2(e13270)* mutants serving as positive controls. All RP mutants showed significantly more resistance to acute oxidative stress (p<0.001) compared to wild-type controls. *rpl-5(0)/+; skn-1(zj15)* double mutant animals were significantly less resistant to oxidative stress compared to *rpl-5(0)/* mutants (p=0.016, third plot). Conversely, *rpl-5(0)/+; daf-16(mu86)* animals displayed similar responses to oxidative stress as *rpl-5(0)/*+ animals (p=0.51, fourth plot). “n” indicates the total number of animals used in the study, all experiments were performed in three biological replicates and all animals were grown at 16°C. *rpl-33(0)/*+^3^ in green, *rpl-5(0)/*+^3^ in orange, *rps-10(0)/+*^1^ in blue, and *rps-23(0)/*+^2^ in red. Balancer chromosomes are denoted as follows: +^1^ = *tmC20*, +^2^ = *tmC5*, +^3^ = *mIn1*. For statistical analysis, Kaplan-Meier analysis with Bonferroni correction was performed for multiple comparisons. **(D)** RNA expression changes of SKN-1 targets in all RP mutants were plotted ^61^.

In acute survival assays using high doses of paraquat, we observed that all RP haploinsufficient strains exhibited significantly enhanced resistance to oxidative stress compared to wild-type controls (**Figure 2C**, first and second plot, p<0.01, Kaplan Meier test with Bonferroni correction). This widespread increase in stress resistance suggests a robust, adaptive mechanism that compensates for elevated reactive oxygen species (ROS) levels.

Perturbations in ribosome biogenesis have been shown to elicit proteotoxicity^49^. Moreover, ribosomal protein haploinsufficiency reduces ribosome levels^50^, which could lead to a decrease in overall protein synthesis. To determine whether the observed elevated levels of oxidative stress in the mutants were due to proteotoxic stress or were related to a reduction in protein synthesis, we pre-treated wild-type animals with inhibitors targeting key pathways: the proteasome (bortezomib), ribosome biogenesis and the translation regulator TORC1 (rapamycin), and translation elongation (cycloheximide), before assessing survival under acute oxidative stress conditions. These treatments significantly enhanced the stress response of wild-type animals (p<0.05 for each drug, Kaplan-Meier test with Bonferroni correction), supporting the role of these pathways in mediating elevated oxidative stress resistance (**Figure S2A**, left panel). Moreover, the combined use of the inhibitors did not further improve survival rates in wild-type animals (**Figure S2A**, right panel). Finally, none of the treatments altered the survival outcomes of *rpl-5(0)/*+ mutants under oxidative stress (p ≥ 0.4, Kaplan-Meier test with Bonferroni correction), suggesting that *rpl-5(0)/*+ mutants inherently possess an elevated baseline oxidative stress response (**Figure S2B**).

Unfolded protein and oxidative stress responses are mediated through SKN-1, which is orthologous to human NRF2^51–55^. SKN-1 further induces a transcriptional response that results in stress resistance when protein translation is inhibited ^56^. Moreover, TORC1 signaling pathway and rapamycin regulate both SKN-1 and DAF-16, orthologous to human FOXO3^57^. Additionally, DAF-16 is involved in repression of RP genes, serving for resistance to hypoxia resistance^58^. Thus, we hypothesized that SKN-1 and DAF-16 might be regulators of the oxidative stress response observed in RP mutants. To dissect these regulatory pathways, we evaluated the oxidative stress survival of *rpl-5(0)/*+ mutants in combination with mutations in *skn-1* and *daf-16* genes. The *skn-1(zj15)* hypomorphic mutation^59^ in the *rpl-5(0)/*+ mutant background significantly diminished oxidative stress survival (**Figure 2C**, third plot, p=0.02, Kaplan-Meier test), suggesting that SKN-1 is essential for the resistance of oxidative stress in *rpl-5(0)/*+ mutants. This was further evident by the lack of significant survival change in *rpl-5(0)/*+ mutants when crossed with the *daf-16(mu86)* mutation^60^, highlighting SKN-1’s unique contribution (**Figure 2C**, fourth plot, p=0.5, Kaplan-Meier test). We observed a mild overexpression of genes upregulated in response to *skn-1* gain of function across the RNA-seq dataset for RP haploinsufficient mutants (**Figure 2D**^61^). This finding suggests that SKN-1 activity is not limited to *rpl-5(0)/*+ mutants but extends to other RP mutants as well.

In *S. cerevisiae*, aberrations in ribosome biogenesis lead to upregulation of targets of Hsf1, a key regulator of proteotoxic stress response^62,49,63^. Thus, we examined the potential overexpression of HSF-1 target genes in all four RP haploinsufficient mutants, by analyzing genes that exhibit differential expression in response to HSF-1 overexpression^64^. Our findings indicate an overexpression trend of genes regulated by HSF-1 among two mutants, *rpl-33(0)/*+ and *rps-23(0)/*+ (**Figure S2C**). We find that the overexpression of HSF-1 targets did not translate into observable differences in acute heat resistance within a 2-hour period (**Figure S2D**, p>0.05). Interestingly, in *C. elegans*, the responses to oxidative stress and heat stress act in opposition and attenuate each other’s effects ^65^. Taken together, these observations suggest a nuanced role for HSF-1 in modulating the cellular stress response under RP haploinsufficiency, where transcriptional activation of stress response genes may not straightforwardly correlate with enhanced stress tolerance.

### Translational Regulation Ensures Ribosome Stoichiometry Balance Despite Haploinsufficiency

To investigate the impact of single-copy ribosomal protein (RP) gene loss on translation, we used ribosome profiling (Ribo-seq) alongside RNA-seq to analyze stage-matched mutant and control *C. elegans* animals at the L4 developmental stage. This approach allowed us to identify translational efficiency (TE) alterations across four RP mutants (**Data S4**). Notably, genes such as *ccdc-47, ddb-1, F32D1.5, pab-1*, and *rad-50* exhibited significantly decreased TE across all RP mutants (p_adj_<0.05), with *pab-1* and *rad-50* also showing RNA overexpression, hinting at a potential compensatory mechanism in response to reduced TE (**Figure S3A**).

Further analysis revealed distinct expression trends. Cell matrix adhesion and defense response genes were overexpressed at both RNA and TE levels (**Figure 3A**, “RNA and TE over”, **Data S4**, significantly enriched GO category list provided in **Data S5**). In contrast, genes involved in histone H3K36 methylation and sister chromatid segregation were underexpressed at both levels, suggesting a systematic downregulation in these functional categories (**Figure 3A**, “RNA and TE under”, **Data S4, S5**). Given the involvement of H3K36 methylation in crucial processes such as RNA polymerase II-mediated elongation^66^ and the regulation of alternative splicing^67^, a reduction in the expression of components of this pathway could further contribute to alterations in transcription and the diversity of transcript isoforms being produced.

**Figure 3:**
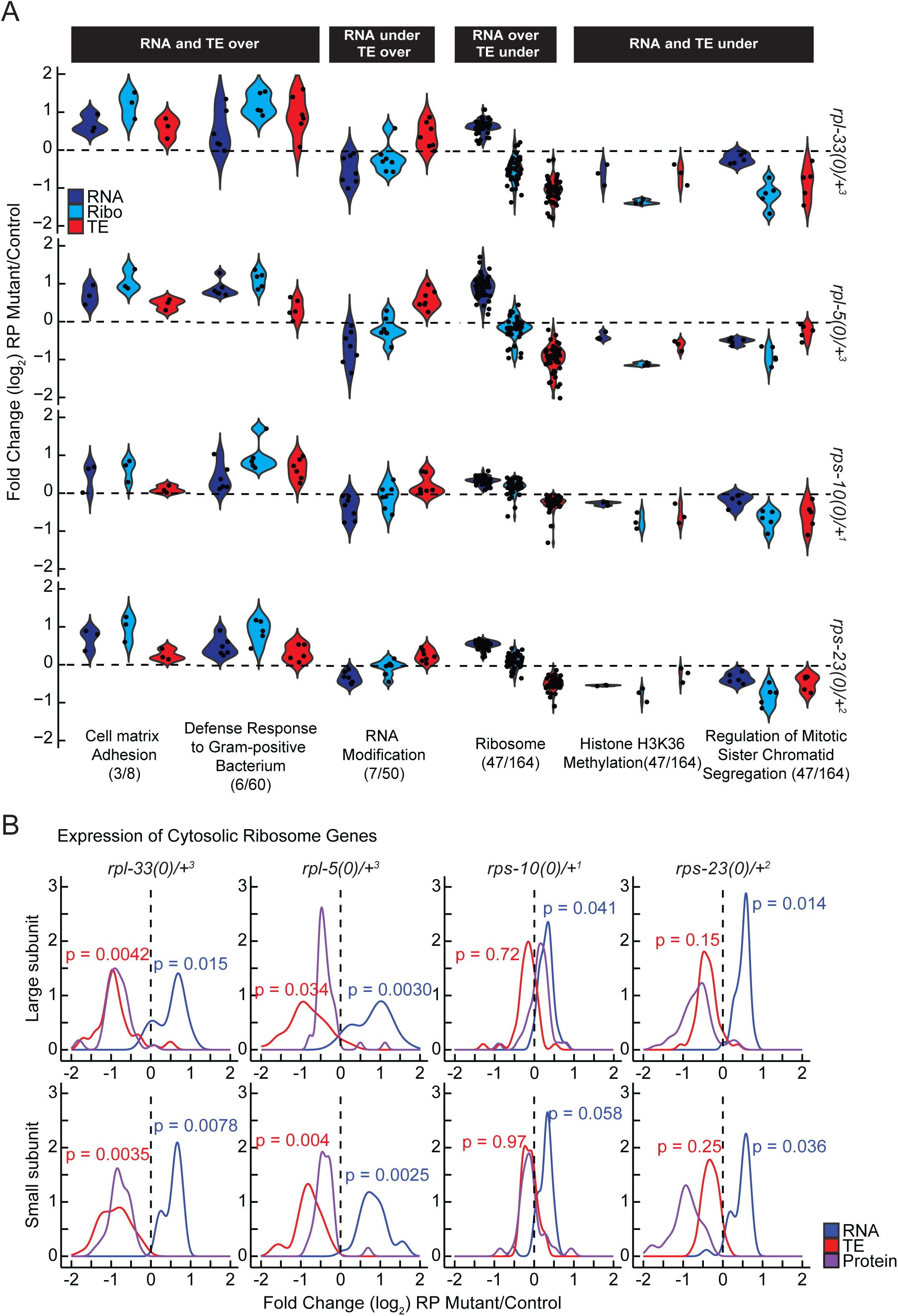
Translational Regulation Ensures Ribosome Stoichiometry Balance Despite Haploinsufficiency. **(A)** Genes were filtered based on collective criteria, including (i) over or underexpression at both the RNA and translation efficiency levels across all mutants, and (ii) altered expression in opposite directions at the RNA or translational efficiency levels, indicating potential buffering effects. Predictions from edgeR were utilized to plot the log_2_ fold changes for selected significant gene annotation (GO) categories, illustrating their RNA levels (blue), Ribo-seq levels (Ribo, cyan), and predicted translation efficiency (TE, red). (**B**) Densities of log_2_ fold changes in RNA, TE, and protein expression of all detected ribosomal protein genes (large or small subunit) in response to the single-copy loss of RP mutants were plotted. The density lines: Blue indicates RNA, red indicates translation efficiency (TE) and purple indicates protein levels.

Interestingly, ribosomal component genes were overexpressed at the RNA level but had reduced TE, notable in both large and small subunit genes (**Figure 3A**, “RNA under TE over” and “RNA over TE under”, respectively, **Data S4**, **S5**). This differential regulation highlights a complex response that balances transcription and translation in response to RP haploinsufficiency to maintain ribosome stoichiometry.

In *S. cerevisiae*, mutations in small subunit genes do not impact the expression of large subunit genes, resulting in an accumulation of unpaired large subunits^68^. In contrast, our observations in *C. elegans* indicate a coordinated response across both subunits at the RNA and TE levels: reductions in ribosomal proteins from either subunit led to a generalized overexpression at the RNA levels (p ≤ 0.01 except *rps-10(0)/+,* ROAST–rotation gene set testing^69^) but a decrease in TE (p ≤ 0.05 for *rpl-33(0)+* and *rpl-5(0)/+,* ROAST) (**Figure 3B**, **S3B**). Given that such a response was not observed in yeast, we asked whether a similar response occurs in response to DBA-specific RP gene reductions in human hematopoietic cells ^50^. Re-analyzing the RNA-seq and Ribo-seq datasets from this study revealed that the results in *C. elegans* mirror those from human *RPL5* and *RPS19* knockdowns in hematopoietic cells, where TE of all subunits are significantly decreased (**Figure S3C**, p<0.05, ROAST, **Data S6**). Hence, the human cellular response is more similar to that in *C. elegans,* in comparison to yeast, highlighting a potentially conserved regulatory mechanism regulating translation for maintaining ribosomal protein stoichiometry under conditions of RP haploinsufficiency.

### Mitochondrial Translation and Morphology Differences in Response to Ribosomal Protein Haploinsufficiency

In RP haploinsufficient mutants, we observed upregulation of glutathione transferase activity and potential activation of SKN-1, a critical factor in maintaining cellular redox balance and facilitating mitochondrial retrograde signaling^70,71^. These observations suggest a potential impairment in mitochondrial function. In particular, genes of the electron transport chain (ETC) need to be translated in a coordinated manner by both cytoplasmic and mitochondrial ribosomes creating a vulnerability in proteostasis^72^.

We next compared changes in RNA expression and TE of ETC components across RP mutants. While we did not observe significant changes in the expression of nuclear-encoded or mitochondrially-encoded ETC components at the RNA level (**Figure-4A**, left, p>0.1, ROAST); a distinct pattern emerged at the TE level. Nuclear-encoded components remained unaffected (p>0.4), whereas mitochondrial-encoded ETC components showed reduced TE across all mutants (**Figure-4A**, right, p<0.005 for *rpl-33(0)/*+ and *rpl-5(0)/*+ and p ≤ 0.16 for *rps-10(0)/*+ and *rps-23(0)/+,* ROAST). This discrepancy suggests that a compensatory mechanism might maintain the stoichiometry of the mitochondrial ETC in response to reduced cytoplasmic ribosome abundance. Although proteomics-level measurements lacked sufficient coverage to quantify corresponding changes in protein abundance for ETC components, the mitochondrially encoded Complex-I component, NDUO-5, was notably underexpressed in *rps-10(0)/*+ animals (∼70% reduction, p_adj_ = 0.2, **Figure-S4A, Data S1**).

Given the marked reduction in the TE of mitochondrial-encoded ETC components across all mutants, we investigated if there was a corresponding change in the abundance of mitochondrial ribosomes. Mitochondrial ribosomal protein mRNAs were generally mildly reduced (p-values < 0.05 for *rpl-33(0)/+, rpl-5(0)/+ and rps-23(0)/+;* p <0.2 for *rps-10(0)/*+, ROAST), however no consistent changes were observed at the TE levels (**Figure S4B**, p-values > 0.5 for all mutants). Similarly, changes at the protein level were not consistent with a wider distribution of mitochondrial ribosomal proteins in *rps-23(0)/*+animals (**Figure S4B**). These results overall suggest that the observed translational efficiency variations across ETC components may not solely be due to differences in mitochondrial ribosome abundance.

The trend of elevated expression of *gst* genes and SKN-1-mediated oxidative stress regulation, both indicative of increased ROS, alongside reduced TE of mitochondrially-encoded ETC components in RP mutants, pointed towards potential mitochondrial dysfunction. Previous studies on mitochondrial dynamics established a link between mitochondrial dysfunction and morphological changes, particularly under stress^73^. Specifically, fission-induced mitochondrial fragmentation, characterized by round mitochondria as opposed to networked mitochondria, is associated with increased ROS and elevated oxidative stress^74,75^. Thus, to investigate the impact of RP haploinsufficiency on mitochondrial morphology, we introduced a body-wall-specific nuclear and mitochondrial GFP marker into the backgrounds of RP haploinsufficient mutants^76^. No differences were observed at 16°C up to the L4 stage, but upon transferring the L4 animals to 23°C and imaging by day three of adulthood, we detected partially penetrant increases in mitochondrial fragmentation across all mutants (**Figure 4B**). To quantify fragmentation, we measured the convexity (degree to which shape differs from its convex hull), defect area (area outside of convex hull), and skeleton branch length of each individual mitochondria. Our results reveal that on average, mitochondria from *rps-10(0)/+, rps-23(0)/+,* and *rpl-5(0)/*+ are significantly more convex, or less networked, compared to wild-type control animals (**Figure 4C**, first plot, p<0.001, Student’s *t*-test). Similarly, all four mutants had a significant average decrease in mean defect area and branch length relative to wild-type control animals (**Figure 4C**, second and third plots, p<0.001, Student’s *t*-test). Together, these results suggest that RP mutants have increased mitochondrial fragmentation that may indicate underlying mitochondrial dysfunction that is both temperature-sensitive and age-related. These results suggest that RP mutants struggle to maintain mitochondrial homeostasis, particularly under moderately higher temperatures, pointing towards a vulnerability in their ability to adapt to environmental conditions.

**Figure 4:**
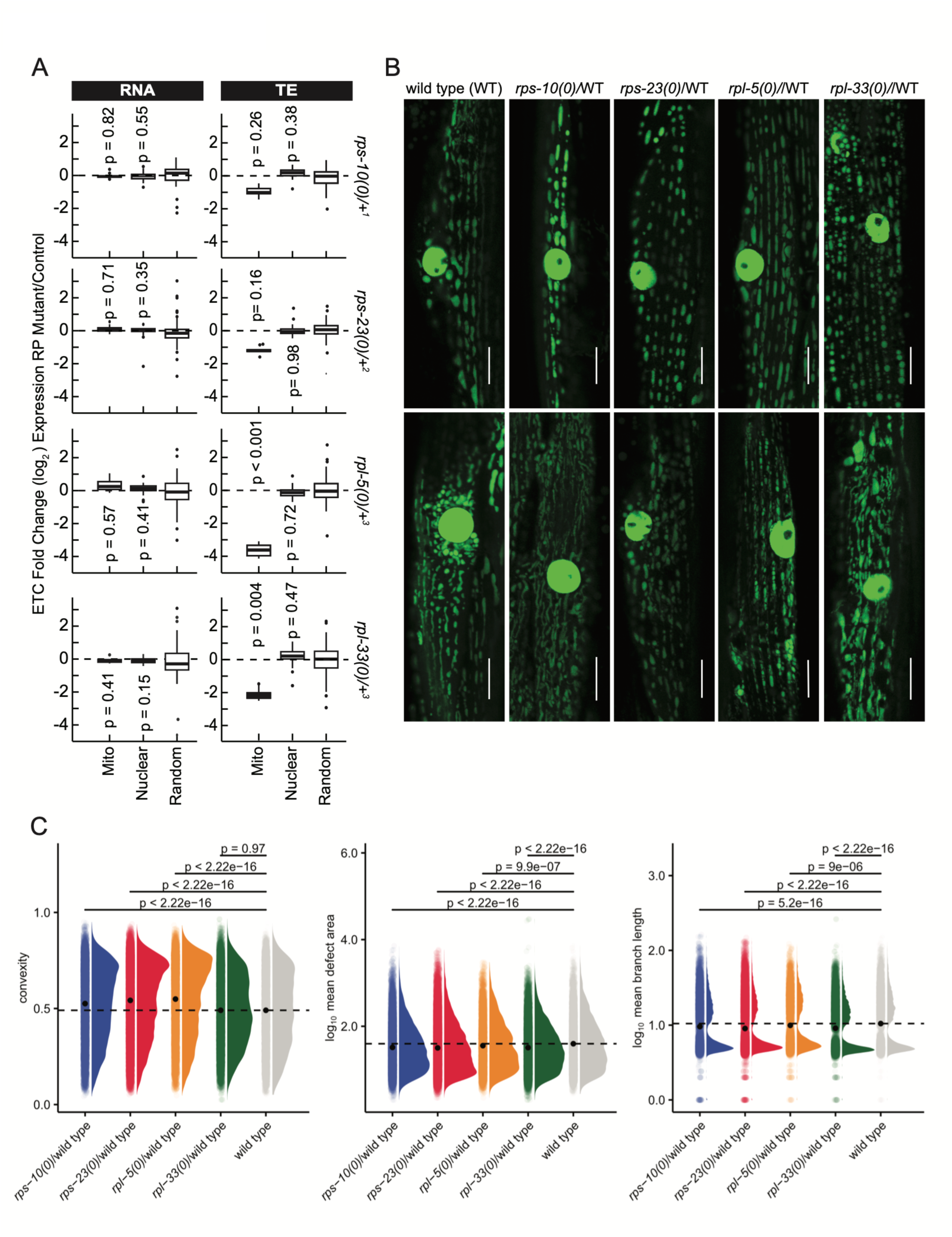
Mitochondrial Translation and Morphology in Response to RP Haploinsufficiency in *C. elegans*. (**A**) RNA expression and translation efficiency differences were plotted along the y-axis for mitochondrial-encoded (mito) or nuclear-encoded (nuclear) genes, as well as for randomly selected genes, for each RP mutant. For statistical analysis of mitochondrial or nuclear-encoded gene expression values, ROAST multivariate gene expression analysis was conducted ^69^. Superscript numbers denote the specific wild type balancer chromosomes, and are used to compare between an RP mutant and the wild-type counterpart. Balancer chromosomes are denoted as follows: +^1^ = *tmC20*, +^2^ = *tmC5*, +^3^ = *mIn1*. (**B**) Representative mitochondrial morphology images using mitochondrial and nuclear localized GFP in body wall muscle cells in RP mutants as well as stage matched wild type control. The images were captured with 63X objective on day three adult animals, which were transferred from 16°C to 23°C on the last two days. RP mutants used here are driven without a balancer chromosome. Images were taken with Stellaris Confocal with a 63X objective, the scale bar represents 10 µm. (**C**) Quantification of the morphology of each individual mitochondria across all mutants and control animals, measured by convexity (degree to which shape differs from its convex hull), defect area (area outside of convex hull), and skeleton branch length. The density distribution of each metric is shown, along with individual mitochondrial measurements. Black dashed line corresponds to the wild type mean. Statistical analyses of the mean of each metric were conducted using an independent Student’s *t*-test relative to the wild-type control. Experiments in B and C were conducted with three biological replicates, analyzing multiple body wall muscle cells from at least 9 animals per group (with >10^3^ mitochondria measurements per strain). Two mitochondria images per mutant and wild type were shown in (B) to represent variability of mitochondrial morphology per body wall cell.

### Mitochondrial Function is Compromised in *rps-10(0)/*+ Mutants

The changes in mitochondrial morphology among RP mutants prompted us to investigate mitochondrial function. Given the link between mitochondrial structure and metabolism^77^, we next evaluated mitochondrial membrane potential using MitoTracker Red CMXRos staining (**Figure S5A**)^78^ and analyzed the overall energy status of the animals by measuring their relative ADP/ATP ratios ^70^.

The *rps-10(0)/*+ mutants exhibited significant decreases in mitochondrial membrane potential, as indicated by significant MitoTracker accumulation (p<0.001, independent Student’s *t*-test) and elevated ADP/ATP ratios (p=0.003, paired Student’s *t*-test), compared to stage-matched controls (**Figures 5A and 5B**). These results reveal that the *rps-10(0)/*+ mutants display a significant disruption in cellular energy homeostasis in addition to compromised mitochondrial membrane potential.

**Figure 5:**
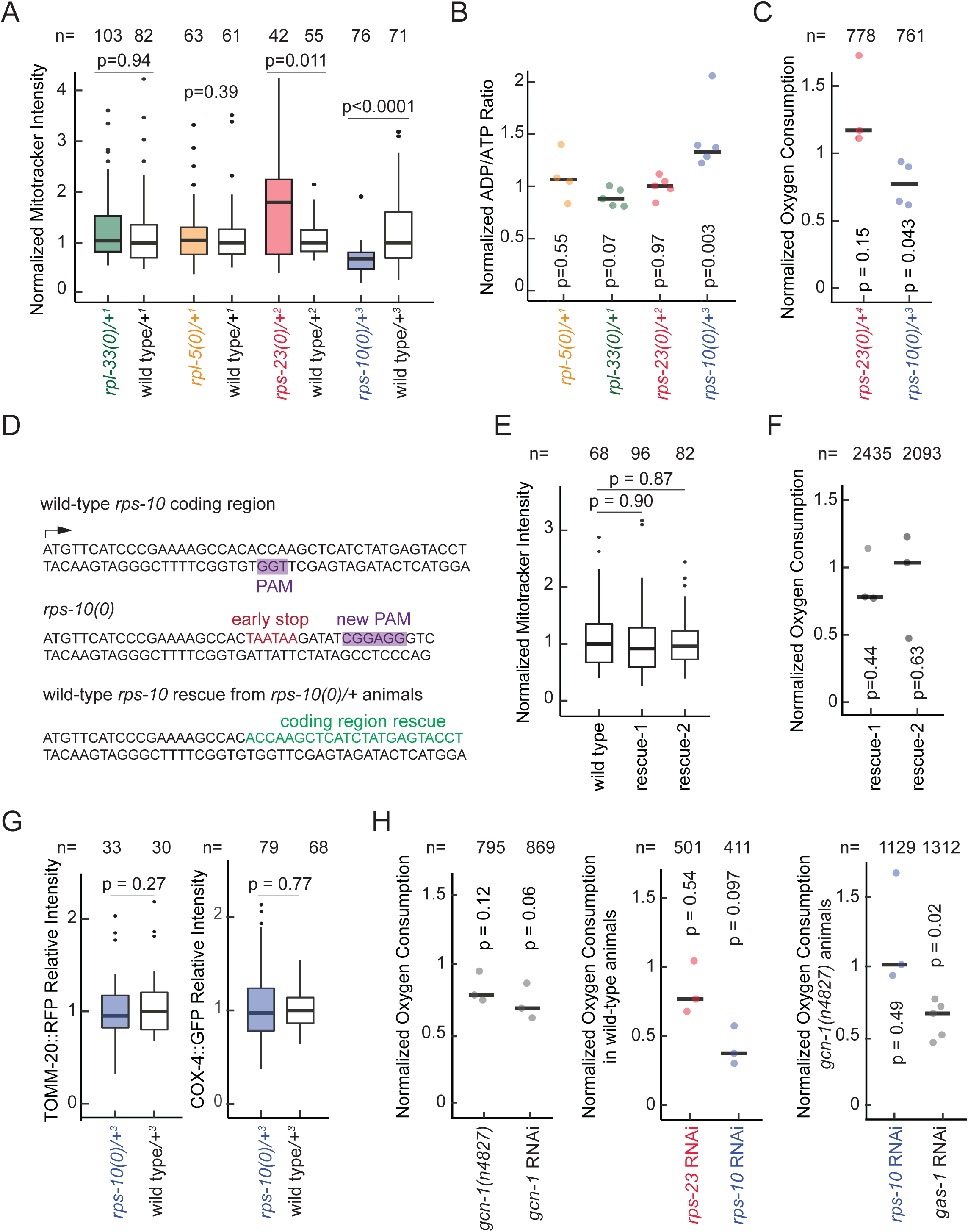
Mitochondrial Function and Energy Levels are Compromised in *rps-10(0)/*+ Mutants. **(A)** Mitochondrial activity, quantified via the uptake of fluorescent MitoTracker Red CMXRos Dye in each RP mutant compared to its wild-type counterpart, was plotted. (**B**) The ADP/ATP ratio, reflecting overall energy levels normalized to stage-matched controls, was presented for each mutant. Measurements from approximately 50 RP mutant and wild-type animals per biological replicate were used, with each replicate represented as a single dot on the plot. (**C**) The average oxygen consumption of *rps-23(0)/*+ and *rps-10(0)/*+ mutants compared to wild type was plotted. Animals were carefully stage-matched to ensure similar body sizes, and oxygen consumption rates were normalized to the number of animals utilized in each assay. (**D**) The genetic modification strategy for the *rps-10* gene is illustrated. The wild-type *rps-10* gene with the PAM sequence highlighted in purple was shown on top; the introduction of tandem early stop codons, generating the *rps-10(0)* variant with a new PAM sequence in purple was indicated in the middle;the restoration of the *rps-10(0)* to its wild-type sequence marked in green was demonstrated at bottom. (**E**) The normalized MitoTracker intensity and (**F**) normalized oxygen consumption were evaluated in wild type and two [*rps-10(0)* to wildtype] rescued strains (rescue-1, rescue-2). (**G**) The relative intensity of TOMM-20::TagRFP and COX-4::GFP levels in *rps-10 (0)/*+ mutants were quantified and plotted against stage-matched wild-type controls. (**H**) Normalized oxygen consumption rates were plotted for *gcn-1(n4827)* mutants and *gcn-1* RNAi compared to their controls (wild type and non-target RNAi) (left). The effects of *rps-10* and *rps-23* RNAi in a wild-type background (middle), alongside the *rps-10* and *gas-1* RNAi effects in *gcn-1(n4827)* mutants (right), with respective controls for each group are shown. All experiments were performed in at least three biological replicates, and the animals were grown at 16°C. For ADP/ATP ratios and oxygen consumption rates, a paired Student’s *t*-test was conducted to account for the coupled nature of these measurements. Mitotracker measurements were analyzed using an independent Student’s *t*-test. Superscript numbers denote the specific balancers compared between an RP mutant and its wild type counterpart. Balancer chromosomes are denoted as follows: +^1^ = *tmC20*, +^2^ = *tmC5*, +^3^ = *mIn1,* +^4^ = *nT1*.

Having observed disruptions in mitochondrial function in *rps-10(0)/*+ but not large subunit RP mutants, we assayed oxygen consumption rates in small subunit RP mutants. *rps-10(0)/*+ mutants showed a reduction in oxygen consumption in comparison to stage matched controls (p<0.05, paired Student’s *t*-test), a trend not observed in *rps-23(0)/*+ (p>0.1, paired Student’s *t*-test) (**Figure 5C**). To ensure decreased oxygen consumption is not due to smaller body size, we quantified the body length and width of *rps-10(0)/*+ animals, relative to control, and found no significant difference between them (**Figure S5B**, width p=0.48, length p=0.08)

Given that energy and mitochondrial functionality changes were specific to the *rps-10(0)/*+ mutants, we sought to determine if they were caused by a background mutation introduced during the CRISPR-mediated early stop integration or possibly due to the maternal inheritance of defective mitochondria. To address these possibilities, *rps-10(0)/*+ hermaphrodites were used to reintroduce the wild-type *rps-10* sequence, leveraging the unique SuperPAM (GGNGG) sequence inserted during the creation of the mutation (**Figure 5D**). This procedure yielded two independent wild-type rescue strains from the F1 generation, each carrying two copies of the reverted wild-type *rps-10* gene. These strains displayed MitoTracker intensities and oxygen consumption rates comparable to control groups, suggesting the observed mitochondrial defects are specific to the single-copy loss of the *rps-10* gene and not related to mitochondrial biogenesis or maternal inheritance (**Figures 5E** and **5F**, p>0.4, independent and paired Student’s *t*-test respectively).

Additionally, we investigated whether mitochondrial defects observed in *rps-10(0)/*+ mutants stemmed from changes in mitochondrial biogenesis or overall abundance. Specifically, we measured mitochondrial DNA levels and observed no differences in mitochondrial DNA content when compared to stage matched wild-type controls (**Figure S5C**). Furthermore, we integrated fluorescent reporters for outer and inner mitochondrial membrane components (TOMM-20::Tag-RFP and COX-4::GFP^79^) into *rps-10(0)/*+ mutants and detected no significant differences in fluorescence intensity compared to stage-matched controls (**Figure 5G**, p>0.3, independent Student’s *t*-test). Taken together, these results indicate that the observed mitochondrial respiration deficits in *rps-10(0)/*+ animals are due to defects in mitochondrial functionality rather than decreased abundance.

Considering the established connections between eIF2α, the integrated stress response (ISR), and mitochondrial function^80–82^, we investigated the role of GCN1, a key factor in the ISR that activates eIF2α kinase GCN2^83,84^. GCN1’s activation of GCN2 requires its interaction with ribosomes^85^, and a cryo-EM study has elucidated that GCN1 binds to collided ribosomes at the interface of the small and large subunits for its role in ribosome quality control (RQC)^86^. Although this structure does not show a physical interaction between RPS10 and GCN1, such a link was implicated through yeast two-hybrid interactions, and correlation between decreased levels of RPS10 and reduced eIF2α phosphorylation, implying a compromised activation of GCN2 ^87^. Decreasing *gcn-1* expression, either through RNAi or a loss-of-function mutation *(n4857)*^88^, resulted in modestly lower oxygen consumption rates (∼ 30% and 20% reduction respectively with p-values 0.12 and 0.06, paired Student’s *t*-test) (**Figure 5H**, left plot). RNAi-mediated knockdown of *rps-10* resulted in approximately a 50% reduction in oxygen consumption in a wild-type background (**Figure 5H**, middle plot, p=0.097). However, in *gcn-1(n4857)* mutants, *rps-10 RNAi* did not exacerbate the decrease in oxygen consumption (**Figure 5H**, right plot, p=0.5, paired Student’s t-test), despite causing developmental delays, thereby validating the effective knockdown of *rps-10*. As a control, we used RNAi against *gas-1*, a gene encoding an ETC component that could lower oxygen consumption even more in the *gcn-1(n4857)* mutants suggesting a floor level of oxygen consumption was not obtained in *gcn-1(n4857)* mutants (**Figure 5H**, right plot, p=0.02, paired Student’s *t*-test). These results overall suggest that (i) disruption in *gcn-1* could lead to reduced mitochondrial function, (ii) in the absence of functional GCN-1, RPS10’s effect on mitochondrial function might be minimized or that a compensatory mechanism is activated.

### Conserved Response to Ribosomal Protein Haploinsufficiency

The similarity of symptoms between DBA and Pearson syndrome^30^, the role of mitochondria in hematopoiesis^29^, and the observed lack of coordination in the expression of mitochondrial components in DBA patients ^31^ led us to re-examine the RNA expression and translation efficiency of genes upon knockdown of the two most frequently mutated RPs (sh*RPS19* and sh*RPL5)* in DBA patients using hematopoietic cells^50^. Specifically, we investigated the conserved orthologs and expression differences in *rpl-5(0)/+ C. elegans* mutants and RPL5 knockdown in human hematopoietic cells (**Data S7**). We particularly wondered if there is conserved unidirectional or bidirectional regulation at the RNA and TE levels impacted by reduced levels of *RPL5* across *C. elegans* and human cells. Similar to our previous results, which suggested translational control to maintain ribosome stoichiometry (**Figures 3B**, **S3C**), we observed significant functional GO term enrichments for both cytoplasmic large and small ribosomal subunits (6.5 and 5.3 fold enrichment, respectively, with p-values < 0.01), characterized by increased RNA levels while TE was decreased, when data from *C. elegans* and humans were combined (Figure S6A, Data S7).

Among unidirectional GO enrichments, we observed ∼2 fold enrichment in the categories of DNA unwinding and mitochondrial matrix among genes that were underexpressed both at the RNA and TE level in *C. elegans* and human cells in response to *RPL5* reduction (**Figure S6A**, **Data S7**). Furthermore, distinct GO categories related to mitochondrial components, especially those associated with the ETC and mitochondrial ribosomes, showed an increase in RNA levels coupled with a decrease in TE (**Figure S6A**), consistent with our results in *C. elegans*. Particularly, significant enrichment of the categories of complex I of the ETC (NADH dehydrogenase activity, **Figure 6A**) and the large subunit of mitochondrial ribosomes (both categories are 2.8 fold enriched, p-values < 0.01, **Figure 6B**) indicate a potentially conserved translational buffering mechanism for mitochondrial ribosomes and ETC components in the face of reductions in cytoplasmic ribosomal machinery.

**Figure 6:**
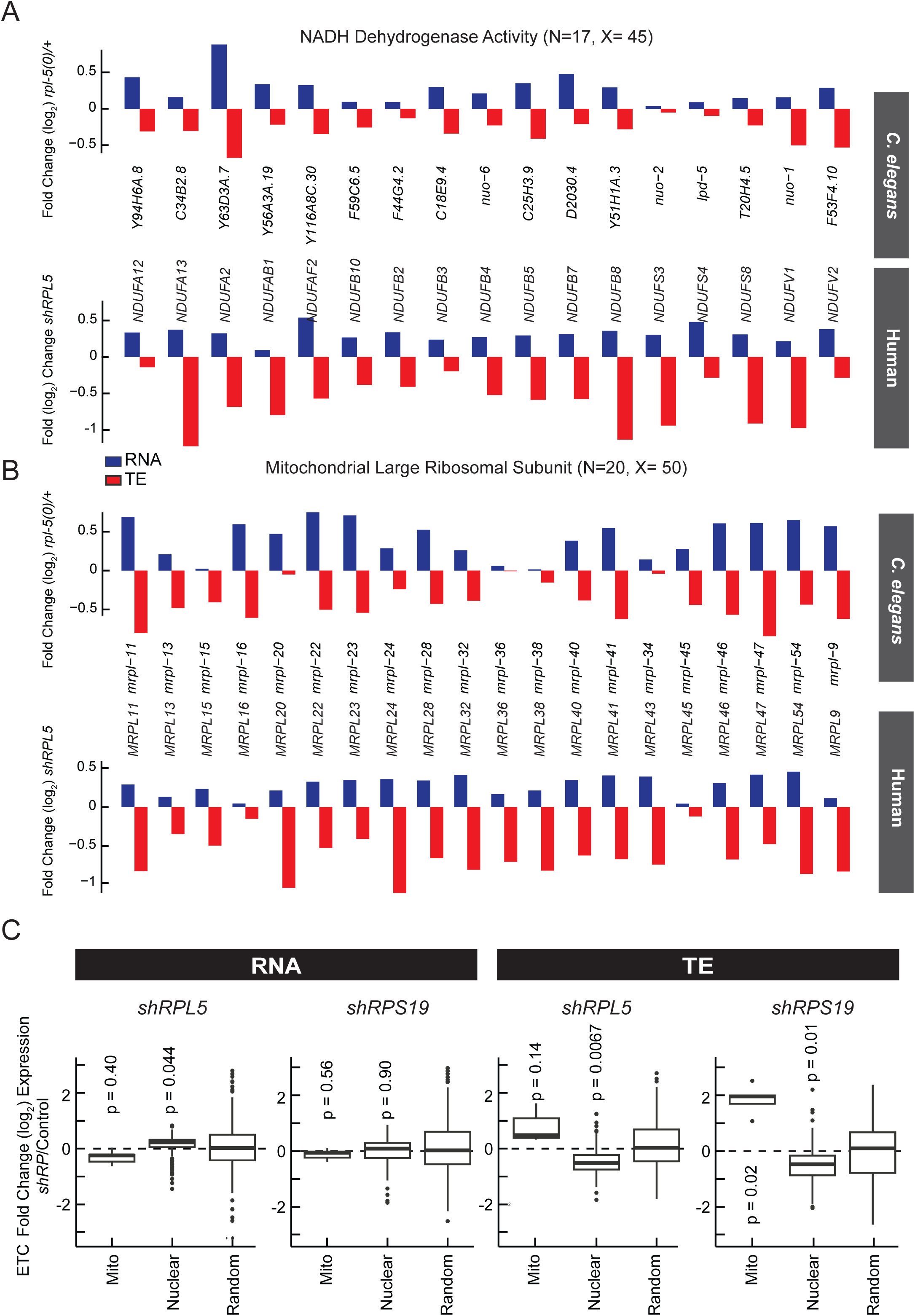
Conserved Translational Regulation in *rpl-5(0)/*+ Animals and *shRPL5* Knockdown in Hematopoietic Cells. Functional gene annotation (GO) enrichment revealed that 17 out of 45 genes within complex I of electron transport chain (the NADH dehydrogenase activity) (**A**) and 20 out of 50 genes associated with the mitochondrial ribosome large subunit (**B**) exhibit similar expression patterns (RNA over, TE under) in both *C. elegans rpl-5(0)/*+ mutants and *shRPL5* knockdown in hematopoietic progenitor cells (p_adj_<0.05, Funcassociate 3.0). (**C**) RNA expression and translation efficiency (TE) differences were plotted along the y-axis for mitochondrial-encoded (mito) or nuclear-encoded (nuclear) electron transport chain (ETC) genes, as well as for randomly selected genes, for *RPL5* and *RPS19* knockdown in blood progenitor cells ^50^. For statistical analysis, ROAST multivariate gene expression analysis was conducted ^69^.

While we observed significantly reduced TE for nuclear ETC components, and both subunits of the mitochondrial ribosome following reductions in *RPS19* and *RPL5* (p < 0.05, ROAST, **Figure 6C**; **Figure S6B**), mitochondrially encoded ETC components were increased at the translation efficiency level (**Figure 6C**, p=0.14, p=0.02, for *RPL5* and *RPS19* reduction respectively, ROAST). These findings overall highlight a broadly consistent effect of *RPL5* reduction on the RNA and TE of critical mitochondrial components in both *C. elegans* and humans, pointing towards a conserved regulatory mechanism.

### Expression Coordination Between Ribosomal and Mitochondrial Components in Human Cells and Impact of RPS10 Reduction on Mitochondrial Activity

To elucidate the gene expression regulatory mechanisms linking mitochondria and ribosomes in human cells, we performed an unbiased co-expression analysis at the transcription and translation levels across lymphoblastoid cells derived from 13 individuals. Specifically, we quantified the similarity of expression patterns across all genes using a compositional proportionality metric^89,90^. This comprehensive analysis unveiled a significant correlation between ribosomal and mitochondrial membrane genes, evidenced by over 1000 significant interactions (**Figure-7A, Data S8**). These findings suggest a highly coordinated regulation of ribosomal and mitochondrial gene expression in human cells (**Figure-S7A**), highlighting the interplay between these essential cellular components.

Building upon these insights into the coordinated expression of ribosomal and mitochondrial genes, we investigated how reductions in cytoplasmic ribosomal proteins affect mitochondrial function, a subject that has been relatively unexplored in human cells. We used K562 leukemia cell line to examine the impacts of reduced levels of specific ribosomal protein transcripts, *RPS10, RPL35A* (the ortholog of *C.elegans rpl-33*), *RPL5,* and *RPS23*. Using siRNA knockdowns, we achieved an approximately 50% reduction in their transcript levels (**Figure S7B**). The mitochondrial membrane potential, assessed using MitoTracker, demonstrated significant decreases in activity following *RPS10* reduction (**Figure 7B**, p = 5e-4, independent Student’s *t*-test), accompanied by changes in the ADP/ATP ratios (**Figure 7C**, p<0.05 for *RPS10* siRNA, paired Student’s *t*-test), highlighting the critical role of ribosomal proteins in supporting mitochondrial energy metabolism.

**Figure 7:**
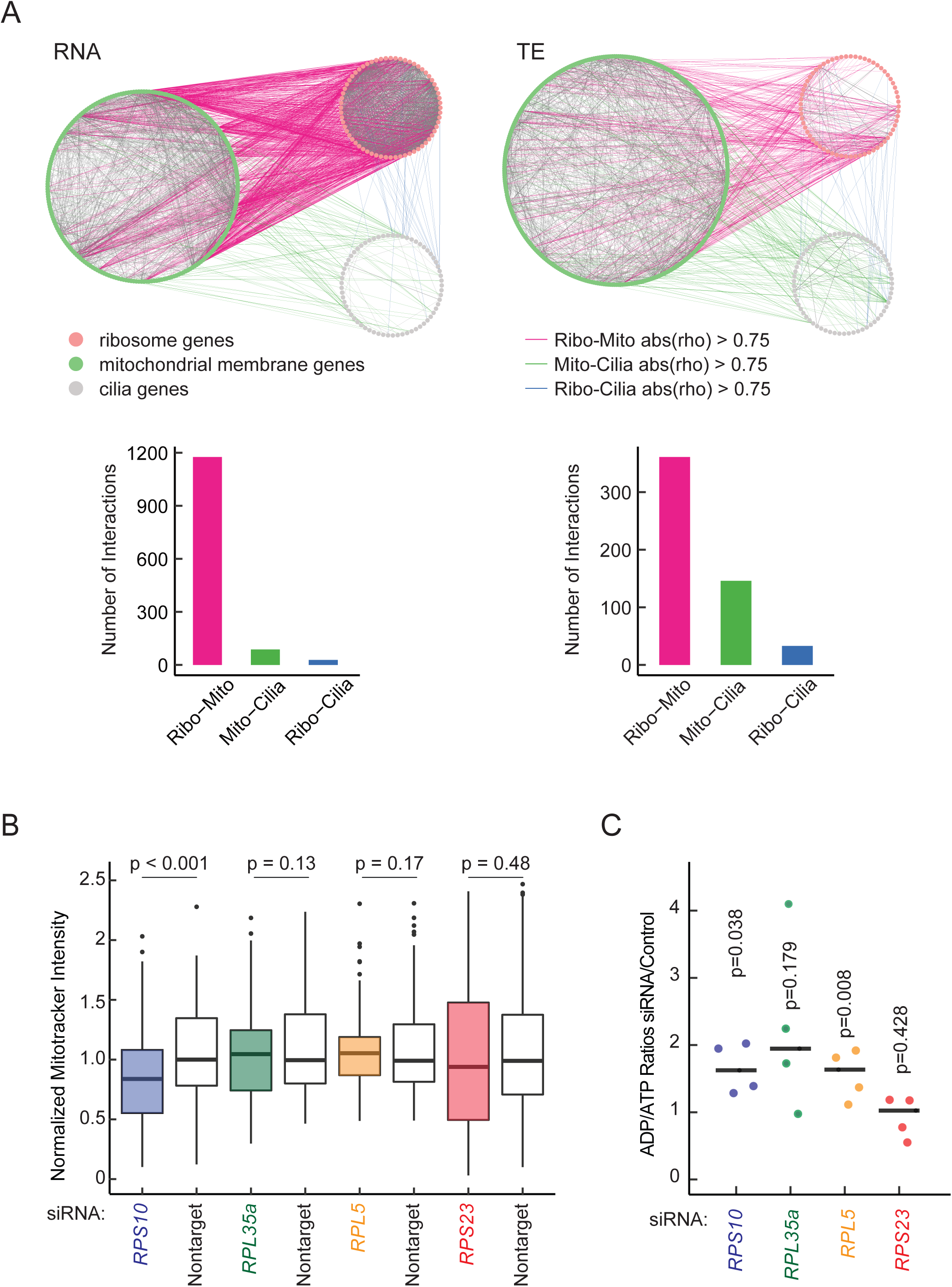
Tight Coordination Between Ribosomal and Mitochondrial Components in Human Cells and Implications of RPS10 Reduction on Mitochondrial Activity. **(A)** Proportionality across all genes was calculated based on 13 human cell lines derived from different individuals ^117^. The significant (rho) interactions between ribosomal and mitochondrial membrane genes at the RNA level (top left) and at the translation efficiency level (TE) (bottom left) were represented by line interaction maps and compared to an unrelated dataset (cilia genes). The number of interactions is displayed at the bottom for RNA and translation efficiency values. Ribosome genes are shown in pink, mitochondrial membrane genes are in green and cilia genes are colored in gray. (**B**) Normalized MitoTracker intensities in response to approximately 50% reduction of *RPS10, RPL35A, RPL5,* and *RPS23* transcripts were plotted in comparison to nontarget siRNA, where mitotracker intensity was quantified for at least 100 cells in three biological replicates, and RNA levels were quantified by quantitative PCR for each replicate (Figure-S7). (**C**) ADP/ATP ratios in response to approximately 50% knockdown of *RPS10, RPL35A, RPL5,* and *RPS23* transcripts were plotted in comparison to non-target siRNA.

These observations collectively emphasize a balance between ribosomal and mitochondrial gene expression, crucial for cellular energy production and metabolic health. The differential regulation of translation efficiency for mitochondrially versus nuclear-encoded ETC components suggests adaptability to counteract the effects of ribosomal protein reduction, with a substantial impact of *RPS10* reduction on mitochondrial activity and energy metabolism.

## DISCUSSION

Here, we investigated the effects of ribosomal protein (RP) haploinsufficiency within the *C. elegans* model focusing on its cellular and developmental consequences. Previous studies have established that functional deficiencies in ribosomal proteins can trigger significant developmental and physiological alterations across a range of model organisms, including *Drosophila* and yeast.^35,40,68,91–94^. In addition to yeast, which possesses widespread genomic duplications of RP genes^95^, or *Drosophila*, which also utilizes specific pathways including the Xrp-Dil8 system activated in response to ribosomal protein knockdown^96,97^, the nematode model provides a unique perspective on the consequences of single copy RP loss.

In this study, we uncovered insights into the relationship between ribosomal stoichiometry, mitochondrial integrity, and cellular metabolism, highlighting the roles of ribosomal proteins beyond their traditional role in protein synthesis. We found that mitochondrial and nuclear encoded components of the electron transport chain are differentially regulated in response to RP reduction or haploinsufficiency in humans and *Caenorhabitis elegans,* respectively. This differential regulation suggests a conserved strategy to maintain mitochondrial function despite variations in cytoplasmic ribosome component expression, suggesting a fundamental connection between ribosomal assembly and metabolic regulation.

Further supporting the tight coupling between ribosomal and mitochondrial components, we discovered significant covariation of these transcripts at both the RNA and translation efficiency levels. At the translation level, this link may be mediated by factors such as TRAP1, which is associated with both mitochondrial and cytoplasmic ribosomes and controls the translation elongation rate^98^. Furthermore, our observations of mitochondrial morphology differences across all RP haploinsufficient mutants, especially those related to RPS10 reduction, indicate substantial impacts on mitochondrial activity in both species. This alteration affects cellular energy homeostasis and suggests that buffering mechanisms for maintenance of mitochondrial health in response to environmental inputs may be compromised.

The mitochondrial metabolism disruptions in *rps-10* haploinsufficient mutants, mirror metabolic changes upon loss of Rpl22A in yeast, such as the decreased one-carbon metabolism, a pathway within mitochondria^91^. Furthermore, assembly of RPS10 in *S. cerevisiae* is disrupted when there is a deficiency of Ltv1, a biogenesis factor. While Ltv1 deficiency provides growth advantage in some conditions, it also predisposes cells to oxidative stress^99^. This raises the question whether the reduction of RPS10 in human leukemia cells and *C. elegans*, leading to mitochondrial dysfunction, is tied to the similar effects seen with reduced Ltv1, given the absence of Ltv1 leads to the absence of RPS10 similar to the effect conferred by the mutation or knockdown.

In addition, previous studies have implicated distinct roles of ribosomal protein paralogs on mitochondrial activity. Specifically, in *Saccharomyces cerevisiae*, only one of the two paralogs of RPL1, RPL2 and RP26 display defective mitochondrial morphology, and RPL1b is involved in translation of respiration related genes^100^. Similarly, in mice, paralog of RPL3, RPL3L, is exclusively expressed in skeletal muscle and heart tissues and its lack is associated with altered ATP levels. Interestingly, RPL3L containing ribosomes interact with mitochondria and potentially interfere with mitochondrial function^101^.

The impact of RP haploinsufficiency on mitochondrial translation efficiency also offers insights into the adaptive mechanisms in response to ribosomal dysfunction. For instance, the overexpression of glutathione transferases observed in *C. elegans* mutants suggests an increased cellular reliance on antioxidant defenses, likely a compensatory response to elevated ROS levels due to mitochondrial dysfunction.

This adaptation reflects an evolutionarily-conserved strategy to safeguard cellular functionality under genetic stress, further emphasizing the interdependence between mitochondrial integrity, ribosomal protein function, and cellular stress responses. Additionally, the regulatory mechanism whereby mitochondrial dysfunction downregulates cytoplasmic protein translation through the phosphorylation of eIF2α by stress-related kinases highlights the reciprocal relationship between mitochondrial stress and cytosolic translation^102,80^. By elucidating the compensatory mechanisms that preserve mitochondrial stoichiometry and function in the face of ribosomal protein loss, we contribute to a deeper understanding of cellular resilience.

## ACKNOWLEDGEMENTS

We sincerely thank Chris Sullivan, and Justin Havird for generously sharing their lab equipment with us. We thank Arlen Johnson, Keiko Torii and Johnson labs for critical discussions. This research is funded by UT Austin, NIH-NIGMS (5R35GM138340) and Welch Foundation (F-2133-20230405).

## MATERIALS AND METHODS

### *C. elegans* Maintenance and Experimental Conditions

*C. elegans* strains used in this study were sourced from the Caenorhabditis Genetics Center (CGC), supported by the NIH Office of Research Infrastructure Programs (P40 OD010440). Standard cultivation practices involved growing the nematodes on Nematode Growth Medium (NGM) plates, which were pre-seeded with the *E. coli OP50* strain. To ensure genetic stability, particularly to minimize recombination and prevent the loss of balancer chromosomes, we housed the animals in an incubator set at 16°C, unless otherwise specified for specific experimental needs.

All haploinsufficient ribosomal protein mutant strains were maintained at 16°C in the presence of wild-type balancer chromosomes, except for the experiments shown in Figure 4B and C. For the *C. elegans* experiments depicted in Figures 1, 2, 3, 4, and 6, the following balancers were used for each haploinsufficient ribosomal protein strain, along with corresponding wild type/balancer controls: r*ps-10(0)/+^1^, rpl-5(0)/+^3^, rpl-33(0)/+^3^,* and *rps-23(0)/+^2^*. For Figure 5C, *rps-23(0)/+^4^* was used, as +^4^ (*nT1)* is homozygous lethal and facilitates the collection of a large sample of animals. For simplicity, Balancers are represented by the “+” symbol followed by a superscript that corresponds to the particular balancer used. Balancer chromosomes are denoted as follows: +^1^ = *tmC20*, +^2^ = *tmC5*, +^3^ = *mIn1*, +^4^ = *nT1*. Balancer chromosomes were used in all control strains, except for the analysis of mitochondrial morphology (Figures 4B and 4C). It’s important to note that we observed consistent genomic instability in RP mutants, characterized by the unusually frequent breakdown of balancer chromosomes when grown at 23°C or at room temperature. Given the critical role of mitochondrial function in maintaining genome stability ^103,104^, we decided to further investigate mitochondrial functionality.

In the study of mitochondrial morphology (Figure-4B and C), initial observations indicated no notable differences in the mitochondrial structures of young adult worms maintained at 16°C. To further explore mitochondrial dynamics under stress, we allowed the animals to mature into three-day-old adults. Subsequently, these adults were subjected to a stress-inducing environment at 23°C for three days prior to the imaging procedures. This temperature shift was implemented to assess potential changes in mitochondrial morphology under slightly elevated temperatures. For Figures 4B and C, the haploinsufficient ribosomal protein (RP) mutants carrying balancers were crossed with males expressing the *myo-3p::GFP* mitochondrial marker. From these crosses, RP haploinsufficient mutant L4 animals without a balancer and carrying a single copy of the *myo-3p::GFP* mitochondrial marker were recovered from the transient F1 generation for use in mitochondrial imaging and analysis.

### Strain Crossing Procedures

#### ESC299 (rpl-5[cc5998,A166X]/mIn1, skn-1(zj15) IV)

Phase 1: Crossed male wild type/*mIn1* (non-dumpy, GFP+) with QV225 (*skn-1(zj15)*) to produce F1 hermaphrodites (wt/mIn1, *skn-1(zj15)/wt)* with GFP. Dumpy F2 progeny were isolated and genotyped for *skn-1(zj15)* homozygosity. Phase 2: Non-dumpy, non-GFP male *rpl-5(0)/*+ were crossed with dumpy, GFP+ hermaphrodites homozygous for *mIn1* and *skn-1(zj15)* from Phase 1. F2 progeny were genotyped for *skn-1(zj15)* homozygosity and *rpl-5(0)* heterozygosity, ensuring all displayed pharyngeal GFP to avoid *rpl-5(0)* homozygous developmental arrest.

#### ESC733 (rps-10[cc2557,T8X], cox-4(zu476[cox-4::eGFP::3XFLAG])/tmC20)

Phase 1: Crossed male *wild-type/tmC20* (non-dumpy, mVenus+) with JJ2586 *(cox-4(zu476[cox-4::eGFP::3XFLAG]))* to select F1 hermaphrodites displaying both body GFP and pharyngeal mVenus. After three generations, dumpy F2 progeny were isolated for homozygosity confirmation in *cox-4::eGFP::3XFLAG* via body GFP+ observation. Phase 2: Non-dumpy, non-GFP male *rps-10(0)/wt* were crossed with GFP+, dumpy hermaphrodites homozygous for *tmC20* and *cox-4(zu476[cox-4::eGFP::3XFLAG])* from Phase 1. F2 progeny were visually genotyped for *COX-4::eGFP* homozygosity and *rps-10(0)* heterozygosity.

#### ESC613 (rps-10[cc2557,T8X]/tmC20, tomm-20::Tag-RFP V)

Phase 1: Crossed male wild-type/tmC20 (non-dumpy, mVenus+) with ESC158 (*tomm-20::Tag-RFP*) to collect F1 hermaphrodites displaying both body RFP and pharyngeal mVenus. Dumpy F2 progeny were isolated and confirmed for tomm-20::Tag-RFP homozygosity via body RFP+ observation. Phase 2: Non-dumpy, non-GFP male *rps-10(0)/*+ were crossed with dumpy, RFP+ hermaphrodites homozygous for tmC20 and tomm-20::Tag-RFP from Phase 1. F2 progeny were visually genotyped for *tomm-20::Tag-RFP* homozygosity and *rps-10(0)* heterozygosity.

For imaging of mitochondria, *C. elegans* with fluorescently labeled mitochondria (*myo-3p::GFP::NLS + myo-3p::mitochondrial GFP*) were crossed into mutant strains along with wild-type worms.

All strains used in this study were provided (**Data S9**).

### Strain Generation via CRISPR-Cas9

Strains ESC 614 and ESC 615 were derived from heterozygous animals carrying the genotype *rps-10[cc2557,T8X]/tmC20, [unc-14(tmIs1219) dpy-5(tm9715)] I.* Young adult heterozygotes were injected with a CRISPR injection mix, which included a 2.5 µM homologous recombination template (ESC-AS-130), 50 ng/µL guide RNA plasmid (pAS14), and 50 ng/µL Cas9-expressing plasmid (pDD132), adapting the co-conversion method^105^. Rescue mutations were initially selected by identifying balancer chromosome, *tmC20*-free adult animals, characterized by the absence of pharyngeal GFP markers and the avoidance of both the developmental arrest associated with homozygous *rps-10(0)* and the uncoordinated phenotype of *tmC20, [unc-14(tmIs1219) dpy-5(tm9715)] I.* progeny. These potential rescue mutations were subsequently confirmed through PCR amplification and Sanger sequencing.

TOMM-20::TagRFP was generated through an in-frame insertion of the TagRFP gene at the C-terminus of the *tomm-20* gene, utilizing a self-excising hygromycin selection-based CRISPR-Cas9 engineering protocol ^106^.

All strains and primer sequences that were generated or used in this study are provided (**Data S19**).

### Proteomics Analysis

*C. elegans* animals were synchronized using a bleach-based solution (0.5M NaOH and 1% NaCl) standard protocol and subsequently grown on NGM plates until they reached the L4 stage. At this point, the animals were collected using a solution of 50 mM NaCl. To ensure the removal of bacteria and prepare for proteomic analysis, the collected animals underwent a series of serial centrifugations at 300 x g for 10 minutes with 3% sucrose in 50 mM NaCl. The animals were then resuspended in Laemmli Buffer (Bio-Rad, 1610737), supplemented with PMSF (Thermo Fisher Scientific, 36978) and BME (Sigma Aldrich, 11411446001), and immediately flash-frozen. Next, the samples were subjected to mechanical disruption via manual bead-beating to ensure thorough digestion of the protein content. The resulting protein fractions were then loaded onto NuPAGE Bis-Tris Gels (4-12%) (Thermo Fisher Scientific, NP0335BOX) and ran briefly using MES SDS Running Buffer (Thermo Fisher Scientific, B0002). The gel was briefly stained by Coomassie Staining (Bio-Rad, #1610786). Post-electrophoresis, the gels within the top stacking portion were cut into sections using a clean razor for further processing. The excised gel sections underwent trypsin digestion before the peptides were desalted. These prepared samples were then analyzed using a Dionex Liquid Chromatography system coupled with an Orbitrap Fusion 2 mass spectrometer. The analytical run was conducted over a 120-minute period to ensure comprehensive peptide identification and quantification.

For data analysis, raw outputs, including Label-Free Quantification (LFQ) values and peptide counts, were processed using Proteome Discoverer version 2.5 ^107^. This software facilitated the mapping of the data against the *C. elegans* reference database. Further quantitative analysis was performed utilizing the DEP (Differential Expression Proteomics) package in R, accessible via the Bioconductor project (https://rdrr.io/bioc/DEP/man/DEP.html). Proteomics analysis by DEP is provided (**Data S1**).

### Body Area Measurement

For the body area and length assays, animals were synchronized through a two-hour egg-laying period and subsequently grown at 16°C until L4 development. The animals were imaged with a 20x Leica SPE DIC microscope. ImageJ software was used to measure the body area by drawing segmented lines along the length of each animal from head to tail (Figure 1D). Body length was quantified by drawing segmented lines from head to tail, and body width was quantified by drawing segmented lines across the midsection of the body (Figure S5D).

### Brood Size Determination

Animals were synchronized using a two-hour egg-laying window. Heterozygous animals, either carrying a ribosomal protein gene mutation and a balancer chromosome or a wild-type gene with the corresponding balancer, were individually transferred to fresh NGM plates. A single animal was moved to a new plate roughly every 24 hours. Hatched progeny from each animal were counted (Figure 1E and S1B). Differences in the mean normalized brood size of each mutant, compared to its respective control, were quantified using an independent Student’s *t*-test.

### Lifespan Assay

The lifespan assay was conducted on solid Nematode Growth Media, both with and without the addition of Fluorodeoxyuridine (FuDR))^108, 109^. Following synchronization via egg-laying, animals were placed on NGM plates with or without 0.5 µM FuDR (Figure S1C and 1F, respectively). The animals were grown at 16°C, and the survival of each nematode was assessed around every 24 hours. Animals on the NGM plates without FUDR were transferred every two days. For FUDR plates, concentrated OP50 spun at 2000 rpm was used as a food source during the lifespan assay. Statistical analysis was conducted using the Kaplan-Meier test with Bonferroni correction for multiple comparisons^110^.

### Acute Oxidative Stress and Heat Stress Assays

For the acute oxidative stress and heat assays (Figure 2C, S2A, S2B and S2D), *C. elegans* were grown at 16°C and synchronized using bleach solution(0.5M NaOH and 1% NaCl). To induce proteotoxicity, translation or TORC1 inhibition, bleached eggs were pre-grown on NGM plates with the following drugs prior to the stress assay: final concentrations of 100 µM rapamycin (MedChemExpress, HY-10219), 25 nM bortezomib (MedChemExpress, HY-10227), and 25 nM cycloheximide (Sigma, 01810-5G).

For the oxidative stress assay, L4 animals were transferred to fresh OP50-seeded NGM plates that had been spiked with 0.2 mM paraquat (Thermo Fisher Scientific, AC227320050). For heat stress, L4 animals were subjected to a temperature of 37°C to induce heat stress.

For both of these assays, the survival rate was evaluated at room temperature across various time points, determining mortality based on the lack of movement within 15 seconds after being gently prodded with a platinum pick. Statistical analysis was conducted using the Kaplan-Meier test with Bonferroni correction for multiple comparisons^110^.

### RNA-seq and Ribo-seq Library Preparation and Data Analysis

For Ribo-seq and RNA-seq procedures, L4 staged heterozygous larvae were collected and remaining bacteria was cleaned up using a 5% sucrose solution with 50 mM NaCl. The animals were stored in 300 µl of lysis buffer (20 mM Tris-HCl pH 7.4, 150 mM NaCl, and 5 mM MgCl2, 1 µl of 1M DTT, 10 µl of 10% Triton-X, 100 ug/ml cycloheximide) and flash frozen in liquid nitrogen. These samples were then stored at −80°C until further use.

The frozen animal pellets were then ground to a fine powder to break cuticles in liquid nitrogen using a mortar and pestle, and the powder was collected in a 1.5 ml tube. The powder was allowed to thaw on ice before 20 units of Turbo DNAse (Thermo Fisher Scientific, AM2238) were added. Each lysate sample was divided into two parts for RNA-seq and Ribo-seq, and 1 ml Trizol (ThermoFisher Scientific, 15596026) was added to the RNAseq aliquot. After a brief vortex and incubation on ice for 15 minutes, the RNA concentration was measured using a Qubit RNA BR assay. RNAse I (Thermo Fisher Scientific, EN0601) was added to each Ribo-seq sample at a ratio of 150 units per 30 µg of RNA and incubated for 30 minutes at room temperature. The RNAse I reaction was stopped with a final concentration of 25 mM ribonucleoside vanadyl complexes (Sigma, R3380). The samples were loaded onto the 34% sucrose cushion prepared in lysis buffer, and spun at 70,000 rpm for 4 hours at 4°C using a TLA 100.3 Rotor in an Optima Ultracentrifuge (Beckman, 361889). After centrifugation, the supernatant was removed to isolate the pellet, which was then dissolved in 1 ml Trizol. The samples with Trizol added for RNA-seq and Ribo-seq were briefly vortexed. Following a 5-minute room temperature incubation, 200 µl of chloroform were added, and the samples were spun at 15,000 rpm for 10 minutes, then the aqueous layer was transferred to a new tube. The final concentration of 50 mM 3 M NaAcetate (pH 5.5), 5 mM MgCl2, and 2 µl Glycoblue coprecipitant (ThermoFisher Scientific, AM9515) was added along with 500 µl isopropanol, and the samples were incubated overnight at −20°C. The next day, the samples were spun at 15,000 rpm, at 4°C for 60 minutes and the pellets were washed with 80% ethanol. The pellets were dissolved in DEPC treated water before the subsequent steps. The Ribo-seq samples were treated with T4 PNK (ThermoFisher Scientific, EK0031) in T4 PNK buffer for 30 minutes at 37°C, and were run on a 15% TBE urea gel. After staining with SYBR Gold (ThermoFisher Scientific, S11494) and imaging, ribosome footprints between 26 and 34 bases were cut out for further processing with D-Plex Small RNA sequencing kit with unique molecular identifiers (UMI) (Diagenode, C05030001, C05030021) for library preparation. For RNA-seq library preparation, SMARTer Stranded RNA-Seq Kit (TakaraBio, 634837) was used. Three independent biological replicates were performed. Each wild type control included two samples that are time-matched and stage-matched to RP mutants to avoid any gene expression changes that are due to the observed developmental delay in RP mutants. Transcriptome mapping reads for the third replicate were not sufficient (less than 300K reads), therefore the third replicate was removed from further analyses.

To analyze human shRPS19 and shRPL5 knockdown experiments in hematopoietic cells, raw data was downloaded from NCBI GE (GSE89183, ^50^).

For read mapping and further processing of the data, Riboflow nextflow pipeline was utilized ^111^(https://github.com/ribosomeprofiling/riboflow). Before mapping the unique identifier barcodes that were added in Ribo-seq, libraries were collapsed to count the number of unique RNA molecules and adapters which were removed. For mapping of reads, *C. elegans* transcriptome (Ensembl, WBCel235) and human transcriptome (Gencode, GRCh38.p14) was used. The Ribo-seq and RNA-seq counts obtained from the pipeline were analyzed using the edgeR pipeline in R ^112^. To investigate differences in translation efficiency (TE), ribosome-bound RNA (Ribo) levels, and RNA expression levels across the samples, three specific contrasts were constructed. Subsequently, a quasi-likelihood F-test (glmQLFTest) was employed to assess these contrasts and adjust for multiple testing errors using a false discovery rate (FDR) approach, setting a p-value threshold of 0.05 for significance. The contrast specifically designed to evaluate translation efficiency was defined as TE_RPmutantvsControl = (RP.Ribo - RP.RNA) - (Control.Ribo - Control.RNA). Here RP indicates a ribosomal protein haploinsufficient strain. This approach allowed for comparison of the TE between ribosomal protein (RP) mutants and their respective controls. When examining significance across all RP mutants, we categorized samples into two groups: RP mutants versus Controls. However, when analyzing individual mutants, each was compared to both stage matched and time matched controls, within the same genetic background.

RNA-seq, Ribo-seq and TE analysis results by EdgeR are provided (**Data S2, S4, S6**).

Mitochondrial genes were extracted using Mitocarta 3.0 ^113^. *C. elegans* and human gene orthologs were extracted using BioMart ^114^. Gene expression values of all orthogonal genes that were conserved between C. elegans and humans are provided (**Data S7**).

### Imaging and Analysis of Mitochondria Morphology

For imaging of mitochondria, *C. elegans* with fluorescently labeled mitochondria (*myo-3p::GFP::NLS + myo-3p::mitochondrial GFP*) were crossed into mutant strains along with wild-type worms. Nematodes were then incubated at 16℃ until the L4 stage was reached. Upon L4 stage development, nematodes were transferred to 23℃ where they were incubated until day three of adulthood. On day three of adulthood, nematodes were immobilized using 10mM Levamisole and imaging of mitochondria was performed using a Leica Stellaris 8 Confocal Microscope (Figure 4B). At least nine nematodes were imaged for each mutant, spanning three biological replicates, with 5-10 cells being imaged in each nematode.

Image analysis was performed using the ilastik v1.4.0 “Pixel Classification + Object Classification” pipeline ^115^. Briefly, raw images were input for pixel classification to separate mitochondria and nuclei from the background. Nuclear localized GFP was excluded for the remaining analyses, and only mitochondria localized GFP was used. The object classification portion of the pipeline was used to extract various features of each individual mitochondria. For each condition more than 10,000 data points were collected. Statistical comparison of mean features, specifically convexity, mean defect area, and mean branch length, between control and mutant samples was carried out using a Student’s *t*-test, performed with the ̀t.test̀ function in R (https://www.R-project.org/). Visualization of feature distributions was performed using the ̀ggplot̀ package in R (Figure 4C, ^116^). The pipeline and code used in this analysis is available at https://github.com/raqmejtru/mito_image_analysis/.

### *C. elegans* Mitotracker Accumulation Assay

The Mitotracker staining of *C. elegans* was adapted from the protocol ^78^. The animals were bleach-synchronized and grown at 16°C until they reached the L4 developmental stage. Subsequently, both the nematodes and OP50 bacteria were collected using M9 buffer, and Mitotracker CMXRos at a concentration of 1 µg/mL (Thermo Fisher Scientific, M7512) was introduced to the samples for staining. The staining procedure involved incubating the animals in Mitotracker solution for 6 hours at 20°C. To remove excess dye, the nematodes were then washed twice with M9 buffer. After washing, the animals were transferred to fresh NGM plates seeded with OP50 and allowed approximately one hour for foraging, which helps in clearing any dye that might have been nonspecifically accumulated in the gut. For quantitative imaging, the stained nematodes were mounted on slides prepared with 3% agarose in M9 buffer, ensuring no anesthetics were used that could potentially interfere with the fluorescence. Imaging was performed using a 20x objective on a Leica SPE microscope using fixed fluorescent exposure. The intensity of the Mitotracker accumulation per area for each animal was quantitatively measured using ImageJ software. Staining specificity was assessed through co-localization studies using a *C. elegans* strain with a CRISPR-engineered knock-in of *cox-4* gene tagged with GFP (*cox-4::GFP*), serving as a marker for mitochondrial inner membranes. These co-localization analyses were conducted using a Leica Stellaris 8 Confocal Microscope with a 63X objective. For visualization and confirmation of the staining pattern’s specificity, we compared the images to the known localization of the COX-4::GFP signal within the mitochondria (**Figure S5A**).

### ADP/ATP Measurement in *C. elegans*

To prepare the samples, *C. elegans* were synchronized using a bleach protocol and subsequently grown on NGM plates seeded with OP50 at 16°C until they reached the L4 developmental stage. Then, the animals were transferred to bacteria-free NGM plates to ensure cleanliness for the assay. A selection of 50 animals was made to specifically exclude those with homozygous balancer chromosomes, and these selected worms were placed in 50 µL of M9 buffer. The prepared samples were flash frozen in liquid nitrogen and stored at −80°C until further analysis. The method for measuring the ADP/ATP ratio in *C. elegans* was adapted from the protocol ^70^. The ADP/ATP ratio was quantitatively measured from lysis supernatant using the Sigma ADP/ATP Assay Kit (Sigma-Aldrich, MAK135-1KT), following the manufacturer’s instructions. Measurements were carried out on a Glomax Luminescence Microplate Reader (Promega).

### *C. elegans* Oxygen Consumption Measurements

To measure oxygen consumption in *C. elegans,* animals were first synchronized using a bleach protocol and subsequently grown on NGM plates that were either seeded with *OP50* or *HT115 E. coli* expressing specific or non-target siRNAs at 16°C until the L4 stage. The animals were collected using a 50 mM NaCl solution and underwent a cleaning process to remove bacteria. This involved centrifugation at 800 x g for 1 minute, repeated three times with M9 buffer to ensure thorough cleansing. After cleaning, the nematodes suspended in the M9 buffer were transferred into the sealed and pre-calibrated chamber of O2k-FluoRespirometer (Oroboros), for the determination of the oxygen consumption rate. The oxygen consumption rate value was then normalized based on the exact number of animals that were introduced into the measurement chamber, to reflect the metabolic rate per individual animal.

### *C. elegans* Fluorescence Intensity Measurements

To measure fluorescence intensity, *C. elegans* were first synchronized through standard bleach protocol and then cultured at 16°C until the L4 stage. The samples were carefully stage matched to control for similar stages and body sizes. The animals expressing TOMM-20::TagRFP and COX-4::GFP, markers for mitochondrial localization, were imaged using a Leica SPE Fluorescence DIC microscope with a 20x objective lens with similar exposure times. For quantifying the fluorescence intensity per animal, the captured images were analyzed using ImageJ software. Specifically, for the measurements of COX-4::GFP fluorescence intensity, care was taken to exclude the pharyngeal area from analysis. This precaution was necessary to avoid interference from the mVenus marker present in both the *rps-10(0)/tmC20* and *wild-type/tmC20* control strains.

### Proportionality Analysis

We conducted a detailed analysis of gene expression samples of 13 individuals from diverse genetic backgrounds. The samples were selected to have Ribo-seq paired with RNA-seq data from the study GSE65912 ^117^, utilizing the RiboFlow toolkit ^111^. To refine the ribosome profiling dataset further, we applied a winsorization technique to adjust for potential PCR duplication artifacts, capping nucleotide counts at the 99.5th percentile to address over-amplified outliers. We excluded 166 human genes identified as lacking polyA tails to ensure the analysis focused on high-quality gene counts. Following this exclusion, both RNA-seq and Ribo-seq data were normalized using counts per million (CPM), selecting genes with a CPM greater than 1 in over 70% of the samples for further analysis. This process resulted in the retention of 10,145 human genes.

TE and RNA value for each gene in each sample was calculated as explained in Ribo-seq analysis section. To assess coordinated expression and TE among genes, we used the ‘lr2rho’ function from the ‘propr’ R package ^89^, inputting CLR values of TE or RNA expression for 10,145 human genes. The resulting rho values, ranging from −1 to 1, facilitated the generation of CoTE and co-expression matrices. Gene sets were curated from the Gene Ontology and KEGG pathway databases, focusing on mitochondrial membrane, ribosomal, and cilia-associated genes. Performing overlap analysis with these gene lists identified 445 mitochondrial, 65 ribosomal, and 124 cilia-associated genes. For comparability, we selected a subset of 65 genes from the cilia-associated gene set to match the ribosomal gene count. The TE and RNA levels of these 575 genes were then analyzed, focusing on interactions where the absolute rho value exceeded 0.75. Gene interactions with a rho value exceeding 0.75 were visualized using Cytoscape version 3.9.1^118^. All Ribo-Mito interactions that had rho values higher than 0.75 are provided (**Data S9**).

### siRNA Knockdown in K562 Cells

K562 Cells (ATCC) were maintained in RPMI (Thermo Fisher Scientific,11879020) + 10% Fetal Bovine Serum/FBS (Gemini Bio, 900-108) + Glucose 200g/L (Thermo Fisher Scientific, A2494001) at 37^0^C + 5%CO_2_. K562 cells were cultured until reaching 80% confluency. The cell density was assessed using a hemocytometer and Trypan Blue (ThermoFisher Scientific, 15-250-061) staining. The cells were centrifuged at 60 x g for 5 minutes, after which the supernatant was replaced with Opti-MEM (Thermo Fisher Scientific, 31985070) to adjust the cell density to 10^6^ cells per mL. siRNA (IDT), at a final concentration of either 12.5 nM or 6.25 nM, was diluted in Opti-MEM and combined with Dharmafect 1 Transfection Agent (Horizon Discovery, NC1308404) at a final concentration of 0.2% v/v. The concentration was pre-determined by initially titrating down siRNA concentrations with an initial qPCR for each siRNA sequence for an approximate 50% reduction in the target RP gene. Transfection Opti-MEM mix was incubated at room temperature for 20 minutes before being added to the cell suspension in Opti-MEM. The cells were then incubated at 37°C with 5% CO2 for 4 hours. After this incubation period, the transfection medium was removed and replaced with RPMI supplemented with FBS and glucose. The cells were harvested 48 hours post-knockdown procedure with details provided in the next section.

After undergoing the knockdown procedure, cells were centrifuged at 600 x g for 5 minutes and subsequently washed with chilled PBS Buffer (Corning 21-040-CM). An aliquot was flash frozen in liquid nitrogen for ADP/ATP assay analysis, and the cells were then subjected to a second centrifugation under the same conditions. The supernatant was discarded, and 350 µL of Trizol was added to the cell pellet. The cells were briefly vortexed before undergoing standard phenol-chloroform precipitation. Next, the cell lysate was treated with Turbo DNase (Thermo Fisher Scientific, AM2238) following the manufacturer’s guidelines and then subjected to acidic phenol-chloroform extraction (ThermoFisher Scientific, AM9720) for RNA purification. The purified RNA was then converted into cDNA using Superscript III Reverse Transcriptase (Thermo Fisher Scientific, 18080-093). Quantitative PCR (qPCR) was performed using the PowerUP SYBR Green Master Mix Solution (ThermoFisher Scientific, A25779) following the manufacturer’s protocol. The qPCR reactions were set up on a Fast 96-well plate (Thermo Fisher Scientific, 4346907) to quantify gene expression levels, allowing for the analysis of the knockdown efficiency and gene expression changes resulting from the RNAi treatment.

Measurement of relative expression was performed using comparative methods through ΔΔCt measurement. Ct (Cycle threshold) values were obtained from qPCR amplification of samples with either their targeted primer or housekeeping primer (GAPDH) in three technical replicates. Average Ct is measured by averaging 3 or at least 2 (only if one technical replication displays value difference more than 0.5) technical replicates. The ΔCt value was calculated by subtracting the average Ct value of a targeted primer with the average Ct value of a housekeeping primer. The ΔΔCt was calculated through subtracting the ΔCt of the treatment group with ΔCt of the control group. The relative expression level was computed with the following equation: Expression level = 2^(-ΔΔCt) ^119^.

siRNA sequences used in this study are provided (**Data S9**).

### K562 ADP/ATP Assay

For the measurement of the ADP/ATP ratio in human cells, following the collection of cells in PBS as described, the ADP/ATP ratio was determined using the ADP/ATP Assay Kit (Sigma-Aldrich, MAK135-1KT). The assay was conducted on a Glomax Luminometer (Promega), following the guidelines provided by the manufacturer.

### K562 Mitotracker Accumulation Assay

In the assay for mitochondrial membrane staining, K562 cells were quantitatively stained using 0.24 µM Mitotracker CMXRos (Thermo Fisher Scientific, M7512) alongside 1 nM Hoechst (Thermo Fisher Scientific, 62249) for nuclear staining, for a duration of 30 minutes. Post-staining, the cells underwent a washing step with warm PBS Buffer, followed by centrifugation at 600 x g for 5 minutes. Subsequently, the cells were resuspended in LiveCell Imaging Solution (Thermo Fisher Scientific, A14291DJ). Fluorescent imaging was performed using a 43x objective on a Leica SPE microscope, and the intensity of Mitotracker staining per area within the cells was quantitatively analyzed using ImageJ software.

## SUPPLEMENTARY FIGURE LEGENDS

**Figure S1: Ribosomal Protein Haploinsufficiency Results in Developmental Delays and Reduced Brood Size Without Affecting Lifespan in *C. elegans***

(**A**) Development of small subunit RP haploinsufficient mutants and their respective wild-type counterparts are shown after 96 hours incubation from embryo at 16℃. Images depict a delay in growth and vulval development. Images taken with differential interference contrast, using a 20X objective. Scale bar represents 50 µm.

(**B**) The brood size of RP mutants and their respective controls were counted and plotted with respect to time. Each line represents a single animal.

(**C**) The lifespan of small subunit RP mutants (top) and large subunit mutants (bottom) were plotted along with their respective controls in the presence of FuDR (5-fluorodeoxyuridine). The Y-axis represents percent survival and “n” represents the total number of animals.

All experiments were done in three biological replicates with animals grown at 16°C. Statistical analysis was performed using Kaplan-Meier test with Bonferroni correction. Superscript numbers denote the specific balancers compared between an RP mutant and its wild type counterpart. Balancer chromosomes are denoted as follows: +^1^ = *tmC20*, +^2^ = *tmC5*, +^3^ = *mIn1,* +^4^ = *nT1*.

**Figure S2: Stress Responses in RP Haploinsufficient Mutants**

**(A)** Survival fractions were analyzed within 8 hours of oxidative stress after pretreatment of rapamycin, bortezomib, and cycloheximide in wild-type animals and *rpl-5(0)/*+ mutants.

(**B**) Wild-type animals were pretreated with double combinations of rapamycin, bortezomib, and cycloheximide and survival fraction within 8 hours of oxidative stress was determined.

(**C**) Density plots of log_2_ RNA expression differences of HSF-1 targets across all RP haploinsufficient mutants were plotted ^64^.

(**D**) Acute Heat Stress Survival: Survival at 37℃ was assessed in all RP mutants to evaluate within 2 hours.

Statistical analyses were performed using Kaplan-Meier test, with Bonferroni correction for multiple samples. All experiments were conducted in 3 biological replicates, all animals were grown at 16°C.

Superscript numbers denote the specific balancers compared between an RP mutant and its wild type counterpart. Balancer chromosomes are denoted as follows: +^1^ = *tmC20*, +^2^ = *tmC5*, +^3^ = *mIn1*.

**Figure S3: Gene Expression and Translation Efficiency (TE) differences in RP Haploinsufficient Mutants**

**(A)** RNA and TE levels for the top 5 most significant genes for all RP haploinsufficient mutants were plotted. The Y-axis shows the log_2_ fold change predictions, with TE and RNA levels for each mutant labeled in distinct colors.

Superscript numbers denote the specific balancers compared between an RP mutant and its wild type counterpart. Balancer chromosomes are denoted as follows: +^1^ = *tmC20*, +^2^ = *tmC5*, +^3^ = *mIn1,* +^4^ = *nT1*.

(**B**) Heatmap depicting the log_2_ fold change of each RP gene expression at the RNA, TE, and protein levels for all four different mutants. The heatmap scale on the left represents non-scaled log_2_ fold changes.

(**C**) RNA expression and TE differences were plotted along the y-axis for all ribosomal protein genes that belong to large subunit (top) and small subunit (bottom) were plotted for *shRPL5* and *shRPS19* knockdown in hematopoietic cells ^50^. For statistical analysis, ROAST multivariate gene expression analysis was conducted ^69^.

**Figure S4: Analysis of Electron Transport Chain (ETC) Component Peptide Counts and Mitochondrial Ribosome Expression in Haploinsufficient RP Mutants**

**(A)** Raw peptide counts from all three replicates were combined to analyze the ETCcomponents, differentiating between those encoded by nuclear genes (red) and those encoded by mitochondrial genes (blue). Although coverage of mitochondrially encoded ETC components was lower, nuclear-encoded components showed a nearly diagonal pattern, indicating minimal deviation from expected levels. The Y-axes of all four graphs display peptide reads from heterozygous ribosomal protein mutants, while the X-axes correspond to stage-matched wild-type controls.

(**B**) Overall, RNA expression, TE, and protein level changes of all mitochondrial RP genes were plotted in RP haploinsufficient mutants. Y-axes show mitochondrial ribosomal proteins that belong to large or small subunits (top and bottom plots respectively) and X-axes show log_2_ fold changes that were predicted by EdgeR for RNA and TE, and by DEP for proteins. For statistical analysis, ROAST multivariate gene expression analysis was conducted ^69^.

**Figure S5: MitoTracker CMXRos staining and Mitochondrial Abundance and DNA Coverage in Haploinsufficient RP Mutants**

**(A)** Staining specificity was assessed through co-localization studies using a *C. elegans* strain with a CRISPR-engineered knock-in of *cox-4* gene tagged with GFP (*cox-4::GFP*), serving as a marker for mitochondrial inner membranes. These co-localization analyses were conducted using a Leica Stellaris Confocal System equipped with a 63X objective. A representative image is shown with MitoTracker CMXRos staining (left), COX::GFP (middle) and merged images (right). Yellow color indicates co-localization of the staining with the mitochondrial inner membrane marker, COX-4.

(**B**) Body length and width were assessed for stage-matched *rps-10(0)/*+ and wild-type controls used in oxygen consumption experiments depicted in Figure 5G.

(**C**) Mitochondrial genome coverage was charted for heterozygous *C. elegans* mutants and stage-matched wild-type controls. The X-axes represent mitochondrial genome positions, and the Y-axes show mitochondrial genome coverage normalized to the nuclear genome. Orange lines indicate heterozygous mutant animals, and blue lines represent stage-matched wild-type controls at the L4 stage.

**Figure S6: Enriched GO Categories Indicative of Translational Control in *C. elegans* and Humans**

**(A)** Log_2_ enrichment (LOD - log of odds ratio) values are plotted for significant gene annotation (GO) categories with LOD > 1 and containing fewer than 300 genes (p values <0.05). The plot displays enriched GO categories that display unidirectional or bidirectional regulation at RNA and TE level in both *C. elegans rpl-5(0)/*+ mutants and in sh*RPL5* knockdown in human hematopoietic progenitor cells (left). All GO enrichment lists are provided in Data S7. Human progenitor data was re-analyzed and retrieved from ^50^. GO enrichment analysis were performed using Funcassociate 3.0^120^.

(**B**) Log_2_ RNA expression and translation efficiency differences were plotted along the y-axis for all mitochondrial ribosomal protein genes that belong to large subunit (top) and small subunit (bottom) were plotted for *shRPL5* and *shRPS19* knockdown in hematopoietic cells ^50^. For statistical analysis, ROAST multivariate gene expression analysis was conducted ^69^.

**Figure S7: Quantification of ribosomal protein gene expression in K562 cells and relative ADP/ATP ratios in response to RP siRNA treatments**

**(A)** The distribution of absolute correlations among a total of 51,465,585 gene pairs across the entire dataset, in contrast with the specific subset of 21,905 pairs involving ribosomal (ribo) and mitochondrial (mito) genes was plotted. Notably, the ribo-mito gene pairs are exclusive combinations of ribosomal genes correlated with mitochondrial genes and vice versa, without including any ribosomal-to-ribosomal or mitochondrial-to-mitochondrial correlations. This plot provides an unrestricted view of the correlation distribution, displaying the entire spectrum of correlations without applying a predefined cutoff (unlike Figure 7A), to fully encapsulate the breadth of gene interactions within the dataset.

(**B**) Normalized transcript levels of target ribosomal protein genes (*RPS10, RPL35A, RPL5,* and *RPS23*) relative to GAPDH were quantified following siRNA treatment in K562 cells. The results were plotted to illustrate the changes in expression for each ribosomal protein gene used in mitotracker intensity measurements shown in Figure 7C.

## Supplementary Tables

**Data S1:** Proteomics analysis of all RP haploinsufficient mutants, including predictions for fold changes and p-values calculated using DEP.

**Data S2:** RNA-seq analysis of all RP haploinsufficient mutants, including predictions for fold changes and p-values calculated using EdgeR.

**Data S3:** Significantly enriched gene annotation categories for genes increased or decreased at the RNA level in all RP haploinsufficient mutants. Analysis was conducted using Funcassociate 3.0 with a provided target and background gene list ^120^.

**Data S4:** RNA, Ribo-Seq, and Translation Efficiency (TE) fold changes and p-values for all RP haploinsufficient mutant animals, analyzed using EdgeR.

**Data S5:** Significantly enriched gene annotation categories for genes increased or decreased at the RNA and TE level, either unidirectionally or bidirectionally, in all RP haploinsufficient mutants. Analysis was conducted using Funcassociate 3.0 with a provided target and background gene list ^120^.

**Data S6:** RNA, Ribo-Seq, and Translation Efficiency (TE) fold changes and p-values in response to *shRPL5* and *shRPS19* knockdown in hematopoietic cells, analyzed using EdgeR. Raw data was obtained from NCBI GEO: GSE89183 ^50^.

**Data S7:** RNA, Ribo-Seq, and Translation Efficiency (TE) fold changes of all orthologous genes between *C. elegans* and humans in response to *RPL5* reduction, analyzed using EdgeR. The orthologous genes were identified using BioMart ^114^. In addition, all significant enriched GO categories are provided in response to unidirectional or bidirectional changes in RNA and TE levels across *C. elegans* and humans in response to *RPL5* reduction^120^.

**Data S8:** All Ribo-Mito interactions with a rho score higher than 0.75 at the RNA and TE level are listed.

**Data S9:** Strains generated or utilized in this study, as well as DNA and RNA sequences used, are provided.

## Notes

### Competing Interest Statement

The authors have declared no competing interest.

### Summary of Updates

I fixed a very small typo on the third page. rps-23 was previously labeled as rps-2 On Figure-2, two of the violin plots were green while they were supposed to be red.

