## Supplementary Figures for "Cytosolic Ribosomal Protein Haploinsufficiency affects Mitochondrial Morphology and Respiration"

Surya, A et al., Figure S1

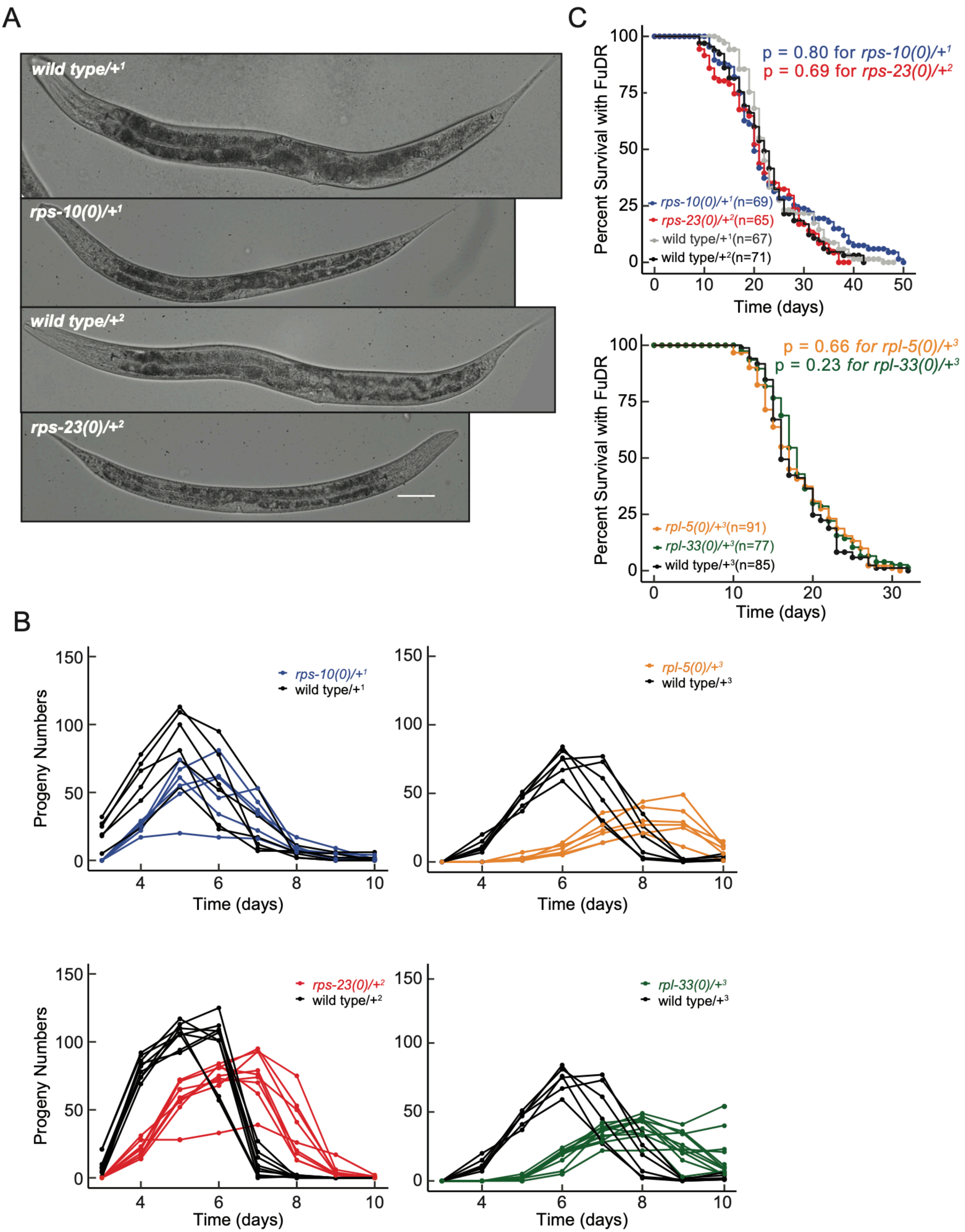

**Figure-S1: Ribosomal Protein Haploinsufficiency Results in Developmental Delays and Reduced Brood Size Without Affecting Lifespan in *C. elegans***

**(A)** Development of small subunit RP haploinsufficient mutants and their respective wild-type counterparts are shown after 96 hours incubation from embryo at 16℃. Images depict a delay in growth and vulval development. Images taken with differential interference contrast, using a 20X objective. Scale bar represents 50 μm.

**(B)** The brood size of RP mutants and their respective controls were counted and plotted with respect to time. Each line represents a single animal.

**(C)** The lifespan of small subunit RP mutants (top) and large subunit mutants (bottom) were plotted along with their respective controls in the presence of FuDR (5-fluorodeoxyuridine*)*. The Y-axis represents percent survival and “n” represents the total number of animals.

All experiments were done in three biological replicates with animals grown at 16°C. Statistical analysis was performed using Kaplan-Meier test with Bonferroni correction. Superscript numbers denote the specific balancers compared between an RP mutant and its wild type counterpart. Balancer chromosomes are denoted as follows: $+^{1}$ = *tmC20*, $+^{2}$ = *tmC5*, $+^{3}$ = *mIn1,* $+^{4}$ = *nT1* .

Surya, A et al., Figure S2

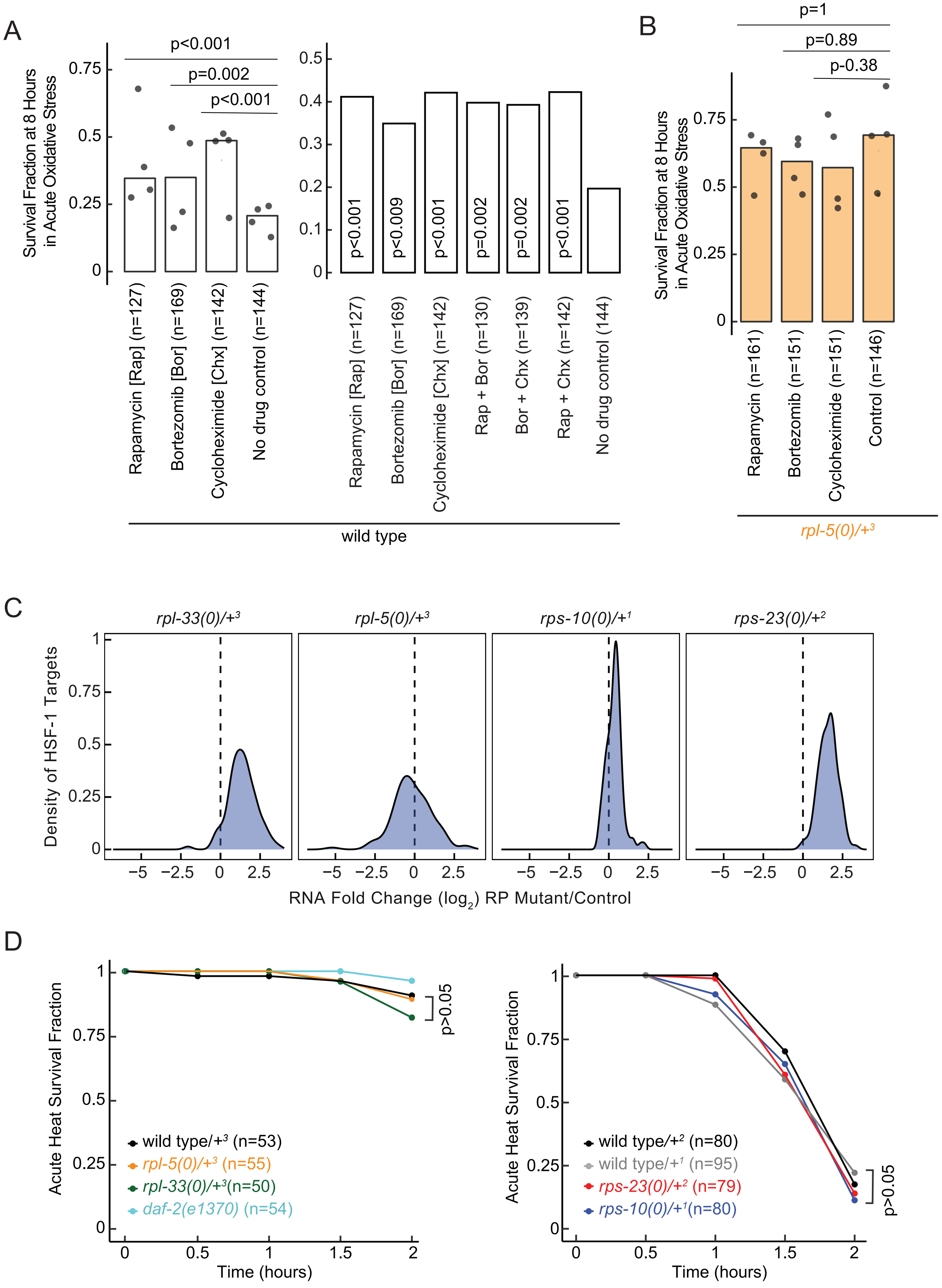

**Figure S2: Stress Responses in RP Haploinsufficient Mutants**

**(A)** Survival fractions were analyzed within 8 hours of oxidative stress after pretreatment of rapamycin, bortezomib, and cycloheximide in wild-type animals and *rpl-5(0)/+* mutants.

**(B)** Wild-type animals were pretreated with double combinations of rapamycin, bortezomib, and cycloheximide and survival fraction within 8 hours of oxidative stress was determined.

**(C)** Density plots of $\log_2$ RNA expression differences of HSF-1 targets across all RP haploinsufficient mutants were plotted [64].

**(D)** Acute Heat Stress Survival: Survival at 37℃ was assessed in all RP mutants to evaluate within 2 hours.

Statistical analyses were performed using Kaplan-Meier test, with Bonferroni correction for multiple samples. All experiments were conducted in 3 biological replicates, all animals were grown at 16°C. Superscript numbers denote the specific balancers compared between an RP mutant and its wild type counterpart. Balancer chromosomes are denoted as follows: $+^1$ = *tmC20*, $+^2$ = *tmC5*, $+^3$ = *mIn1*.

Surya, A et al., Figure S3

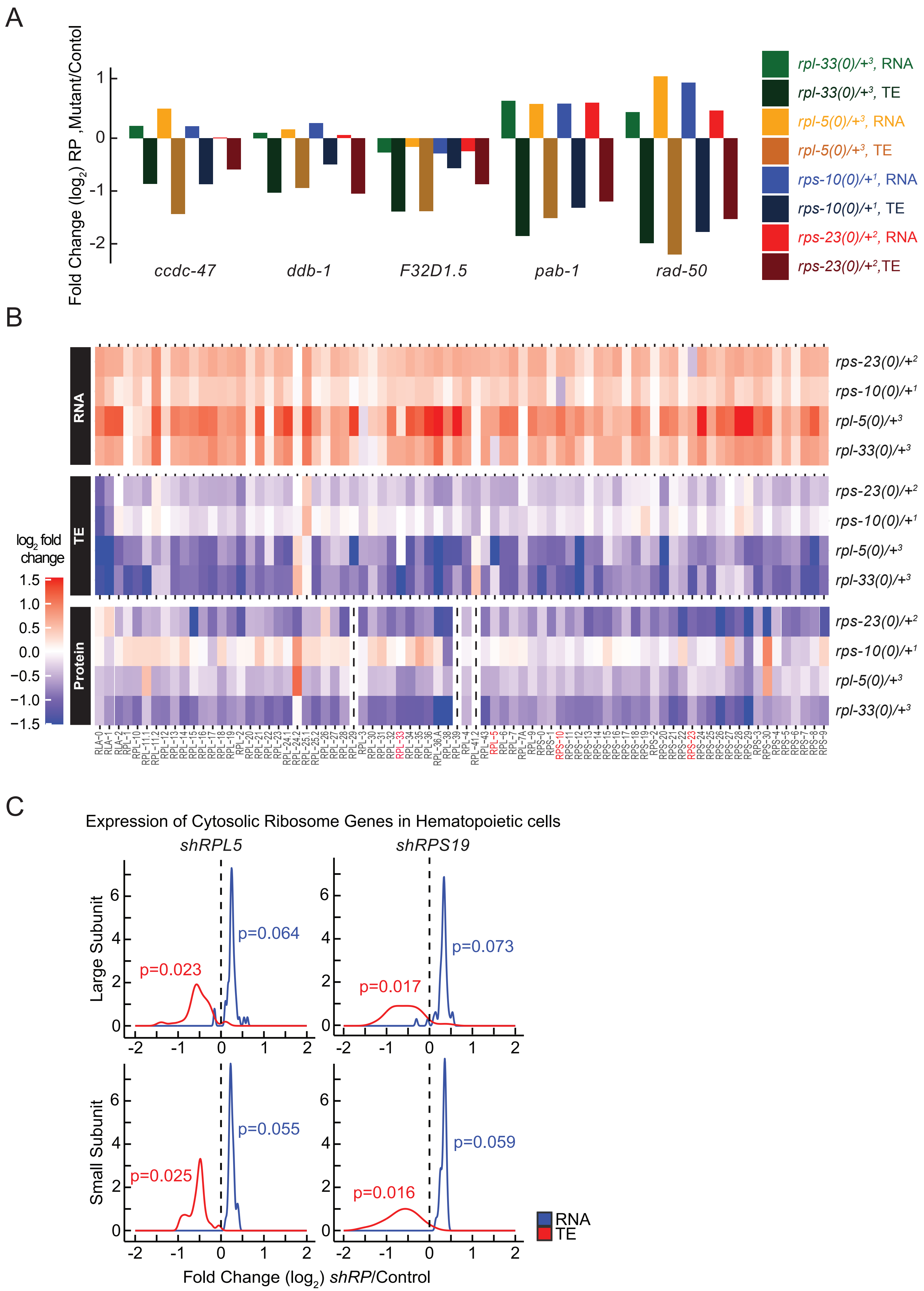

**Figure S3: Gene Expression and Translation Efficiency (TE) differences in RP Haploinsufficient Mutants**

**(A)** RNA and TE levels for the top 5 most significant genes for all RP haploinsufficient mutants were plotted. The Y-axis shows the $\log_2$ fold change predictions, with TE and RNA levels for each mutant labeled in distinct colors.

Superscript numbers denote the specific balancers compared between an RP mutant and its wild type counterpart. Balancer chromosomes are denoted as follows: $+^1$ = *tmC20*, $+^2$ = *tmC5*, $+^3$ = *mIn1,* $+^4$ = *nT1* .

**(B)** Heatmap depicting the $\log_2$ fold change of each RP gene expression at the RNA, TE, and protein levels for all four different mutants. The heatmap scale on the left represents non-scaled $\log_2$ fold changes.

**(C)** RNA expression and TE differences were plotted along the y-axis for all ribosomal protein genes that belong to large subunit (top) and small subunit (bottom) were plotted for *shRPL5* and *shRPS19* knockdown in hematopoietic cells [50]. For statistical analysis, ROAST multivariate gene expression analysis was conducted [69].

Surya, A et al., Figure S4

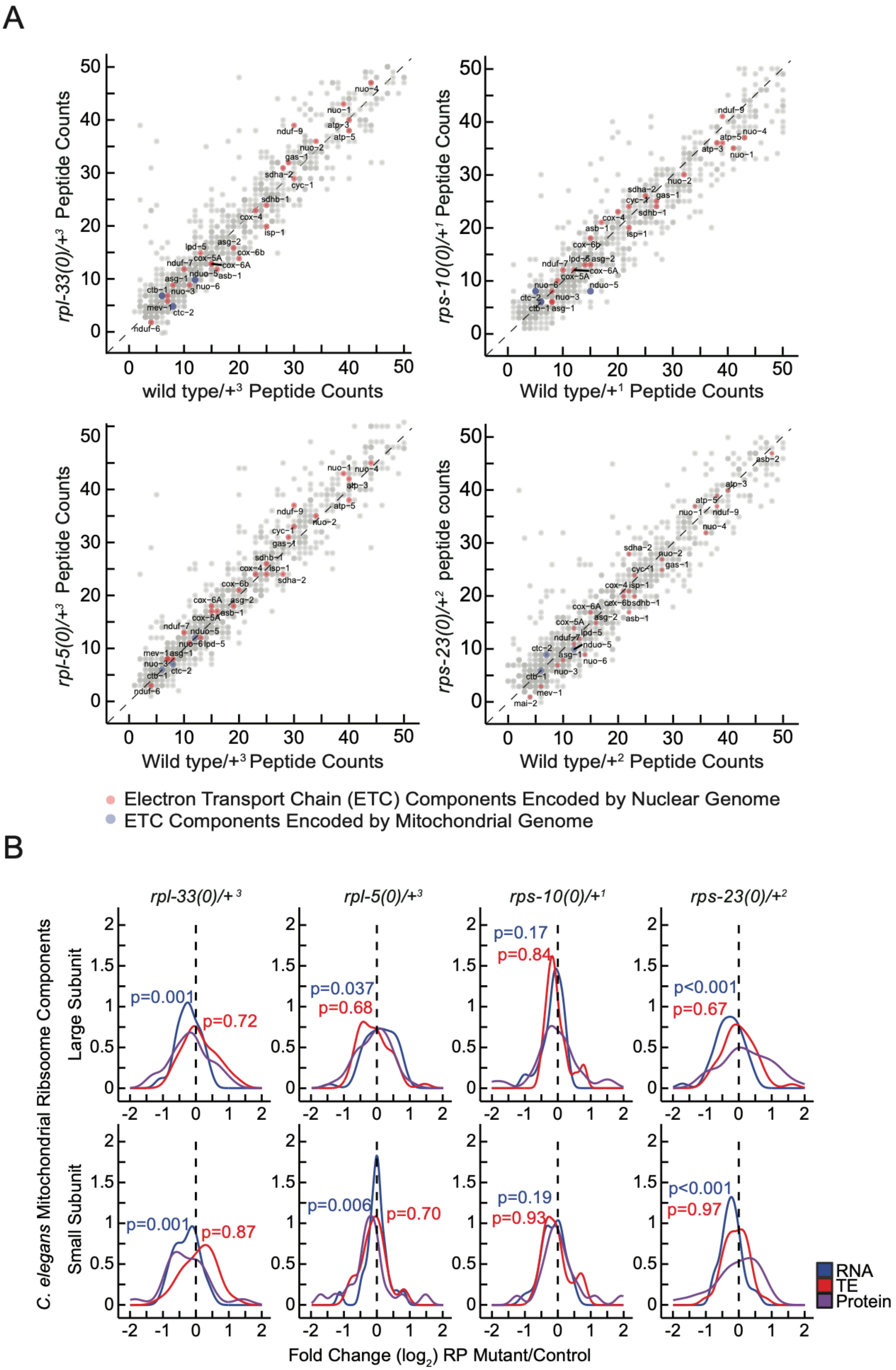

**Figure S4: Analysis of Electron Transport Chain (ETC) Component Peptide Counts and Mitochondrial Ribosome Expression in Haploinsufficient RP Mutants**

**(A)** Raw peptide counts from all three replicates were combined to analyze the ETCcomponents, differentiating between those encoded by nuclear genes (red) and those encoded by mitochondrial genes (blue). Although coverage of mitochondrially encoded ETC components was lower, nuclear-encoded components showed a nearly diagonal pattern, indicating minimal deviation from expected levels. The Y-axes of all four graphs display peptide reads from heterozygous ribosomal protein mutants, while the X-axes correspond to stage-matched wild-type controls.

**(B)** Overall, RNA expression, TE, and protein level changes of all mitochondrial RP genes were plotted in RP haploinsufficient mutants. Y-axes show mitochondrial ribosomal proteins that belong to large or small subunits (top and bottom plots respectively) and X-axes show $log_2$ fold changes that were predicted by EdgeR for RNA and TE, and by DEP for proteins. For statistical analysis, ROAST multivariate gene expression analysis was conducted [69].

Surya, A et al., Figure S5

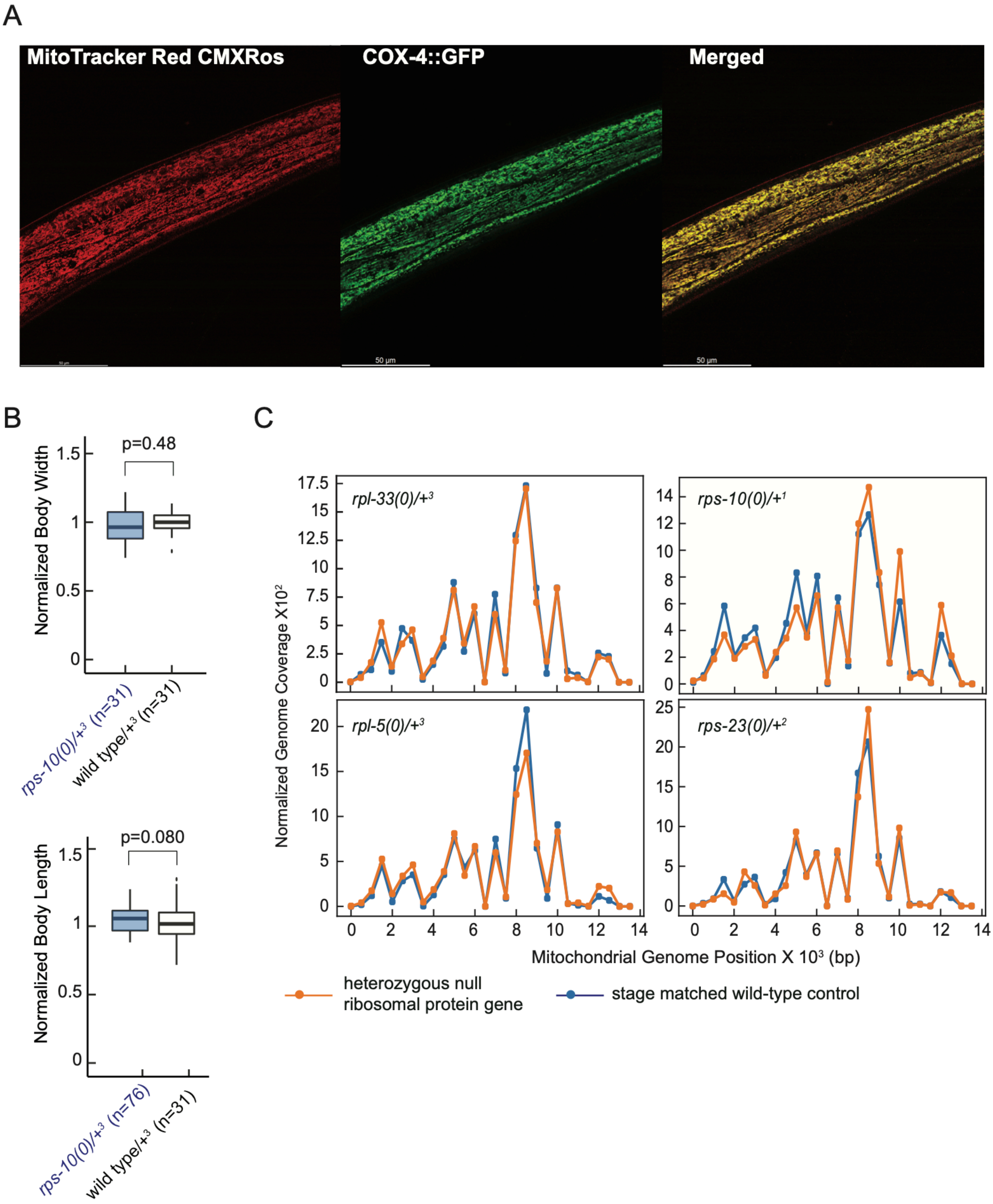

**Figure S5: MitoTracker CMXRos staining and Mitochondrial Abundance and DNA Coverage in Haploinsufficient RP Mutants**

**(A)** Staining specificity was assessed through co-localization studies using a *C. elegans* strain with a CRISPR-engineered knock-in of *cox-4* gene tagged with GFP (*cox-4::GFP*), serving as a marker for mitochondrial inner membranes. These co-localization analyses were conducted using a Leica Stellaris Confocal System equipped with a 63X objective. A representative image is shown with MitoTracker CMXRos staining (left), COX::GFP (middle) and merged images (right). Yellow color indicates co-localization of the staining with the mitochondrial inner membrane marker, COX-4.

**(B)** Body length and width were assessed for stage-matched *rps-10(0)/+* and wild-type controls used in oxygen consumption experiments depicted in Figure 5G.

**(C)** Mitochondrial genome coverage was charted for heterozygous *C. elegans* mutants and stage-matched wild-type controls. The X-axes represent mitochondrial genome positions, and the Y-axes show mitochondrial genome coverage normalized to the nuclear genome. Orange lines indicate heterozygous mutant animals, and blue lines represent stage-matched wild-type controls at the L4 stage.

Surya, A et al., Figure S6

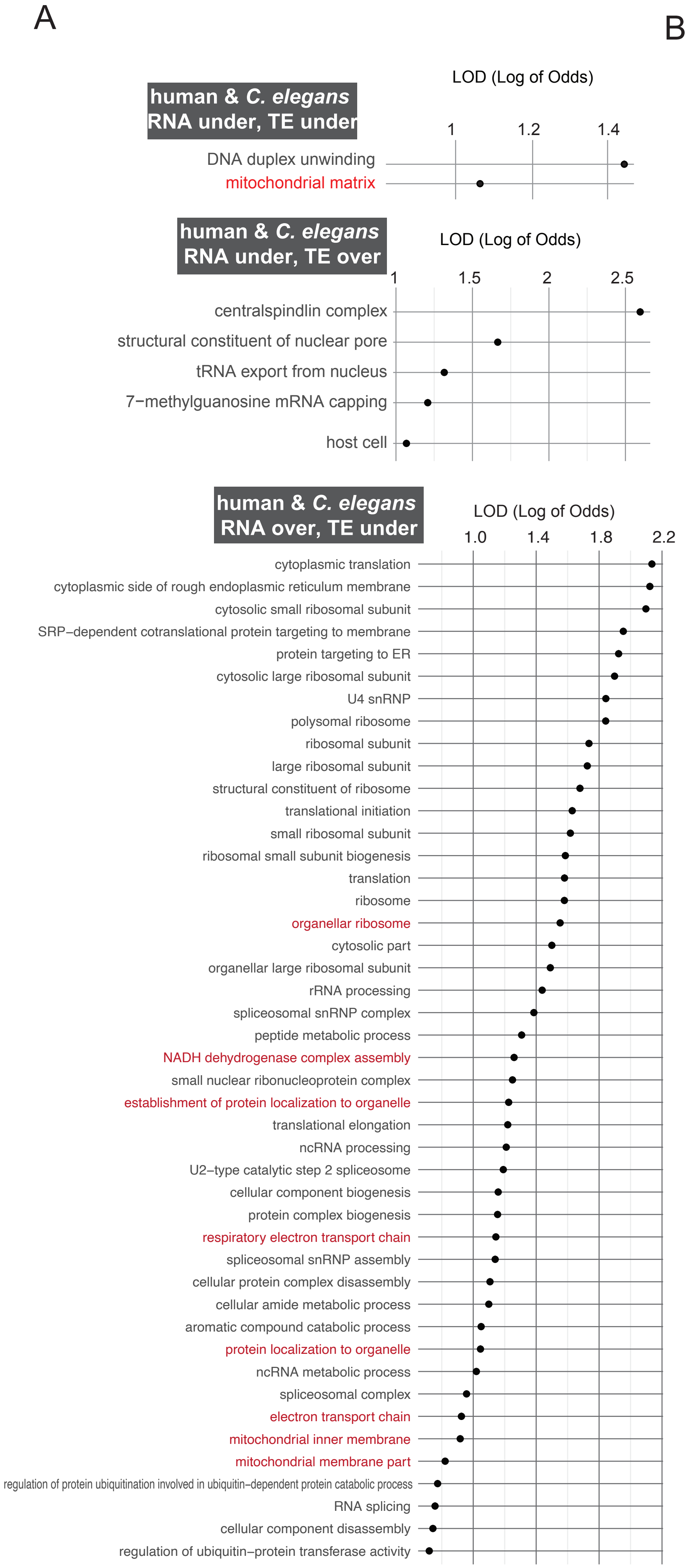


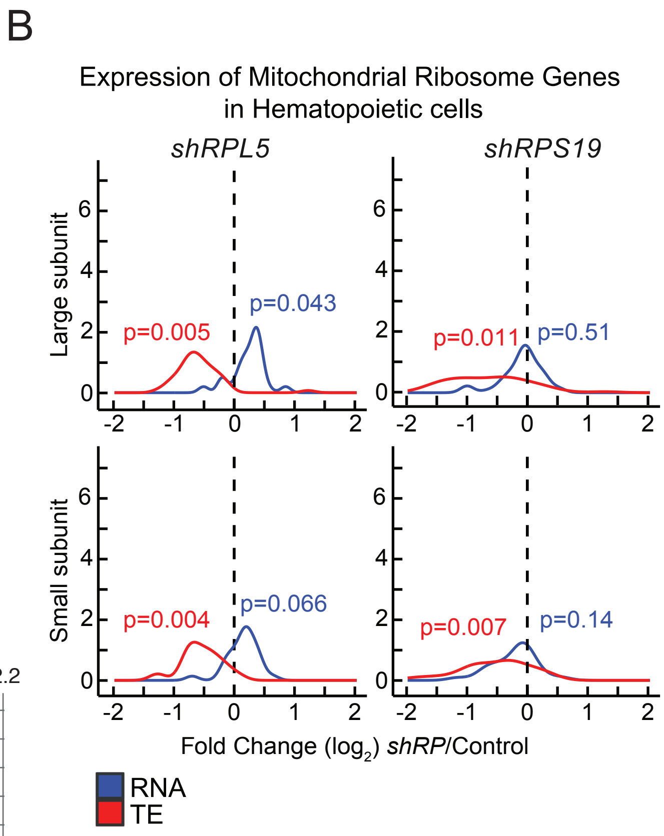

**Figure S6: Enriched GO Categories Indicative of Translational Control in *C. elegans* and Humans**

**(A)** $Log_2$ enrichment (LOD - log of odds ratio) values are plotted for significant gene annotation (GO) categories with LOD > 1 and containing fewer than 300 genes (p values <0.05). The plot displays enriched GO categories that display unidirectional or bidirectional regulation at RNA and TE level in both *C. elegans rpl-5(0)/+* mutants and in sh*RPL5* knockdown in human hematopoietic progenitor cells (left). All GO enrichment lists are provided in Data S7. Human progenitor data was re-analyzed and retrieved from [50]. GO enrichment analysis were performed using Funcassociate 3.0 [120].

**(B)** $Log_2$ RNA expression and translation efficiency differences were plotted along the y-axis for all mitochondrial ribosomal protein genes that belong to large subunit (top) and small subunit (bottom) were plotted for *shRPL5* and *shRPS19* knockdown in hematopoietic cells [50]. For statistical analysis, ROAST multivariate gene expression analysis was conducted [69].

Surya, A et al., Figure S7

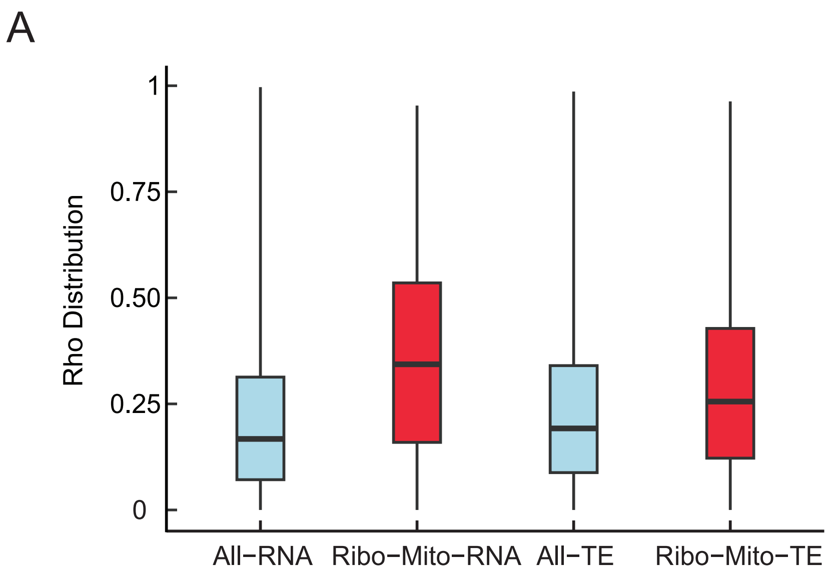


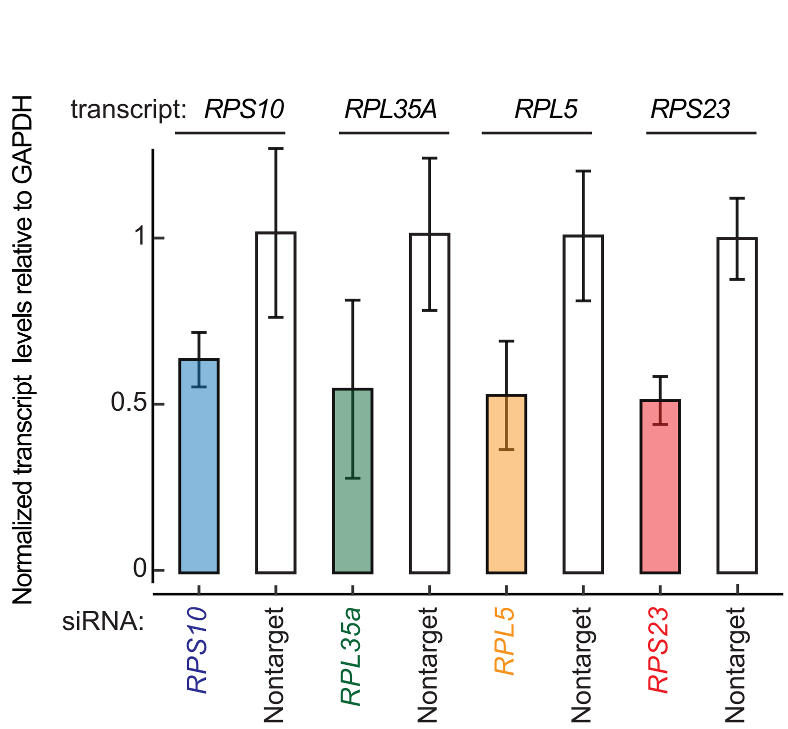

**Figure S7: Quantification of ribosomal protein gene expression in K562 cells and relative ADP/ATP ratios in response to RP siRNA treatments**

**(A)** The distribution of absolute correlations among a total of 51,465,585 gene pairs across the entire dataset, in contrast with the specific subset of 21,905 pairs involving ribosomal (ribo) and mitochondrial (mito) genes was plotted. Notably, the ribo-mito gene pairs are exclusive combinations of ribosomal genes correlated with mitochondrial genes and vice versa, without including any ribosomal-to-ribosomal or mitochondrial-to-mitochondrial correlations. This plot provides an unrestricted view of the correlation distribution, displaying the entire spectrum of correlations without applying a predefined cutoff (unlike Figure 7A), to fully encapsulate the breadth of gene interactions within the dataset.

**(B)** Normalized transcript levels of target ribosomal protein genes (*RPS10, RPL35A, RPL5,* and *RPS23*) relative to GAPDH were quantified following siRNA treatment in K562 cells. The results were plotted to illustrate the changes in expression for each ribosomal protein gene used in mitotracker intensity measurements shown in Figure 7C.
